# A *CDKN2C* retroduplication in Bowhead whales is associated with the evolution of extremely long lifespans and alerted cell cycle dynamics

**DOI:** 10.1101/2022.09.07.506958

**Authors:** Juan M. Vazquez, Morgan Kraft, Vincent J. Lynch

## Abstract

Among the constraints on the evolution of remarkably long lifespans is an increased risk of developing cancer because organisms with long lifespans have more time to accumulate cancer-causing mutations than organisms with shorter lifespans. Indeed, while there is a strong correlation between lifespan and cancer risk within species, there is no correlation between maximum lifespan and cancer risk across species (‘Peto’s Paradox’). Here we use evolutionary genomics and comparative experimental biology to explore the mechanisms by which Bowhead whales (*Balaena mysticetus*), which can live at least 211 years, evolved their extremely long lifespans. We found that the Bowhead whale genome encodes a species-specific retroduplicated *CDKN2C* (p18^INK4C^) gene (*CDKN2CRTG*). The *CDKN2CRTG* gene is embedded within a Cetacean-specific LINE L1 element, and is highly expressed in Bowhead whale tissues likely because it coopted an L1 promoter to drive constitutive expression. Furthermore we use a series of gain of function experiments to show how the duplicate *CDKN2CRTG* gene may influence cellular phenotypes such as cell cycle progression and DNA damage repair in ways that are beneficial for aging and cancer resistance. Remarkably, Bowhead and Right whales only diverged ~4-5 million years ago, suggesting the long lifespan of Bowheads may have evolved relatively recently and coincident with the origin of *CDKN2CRTG*.

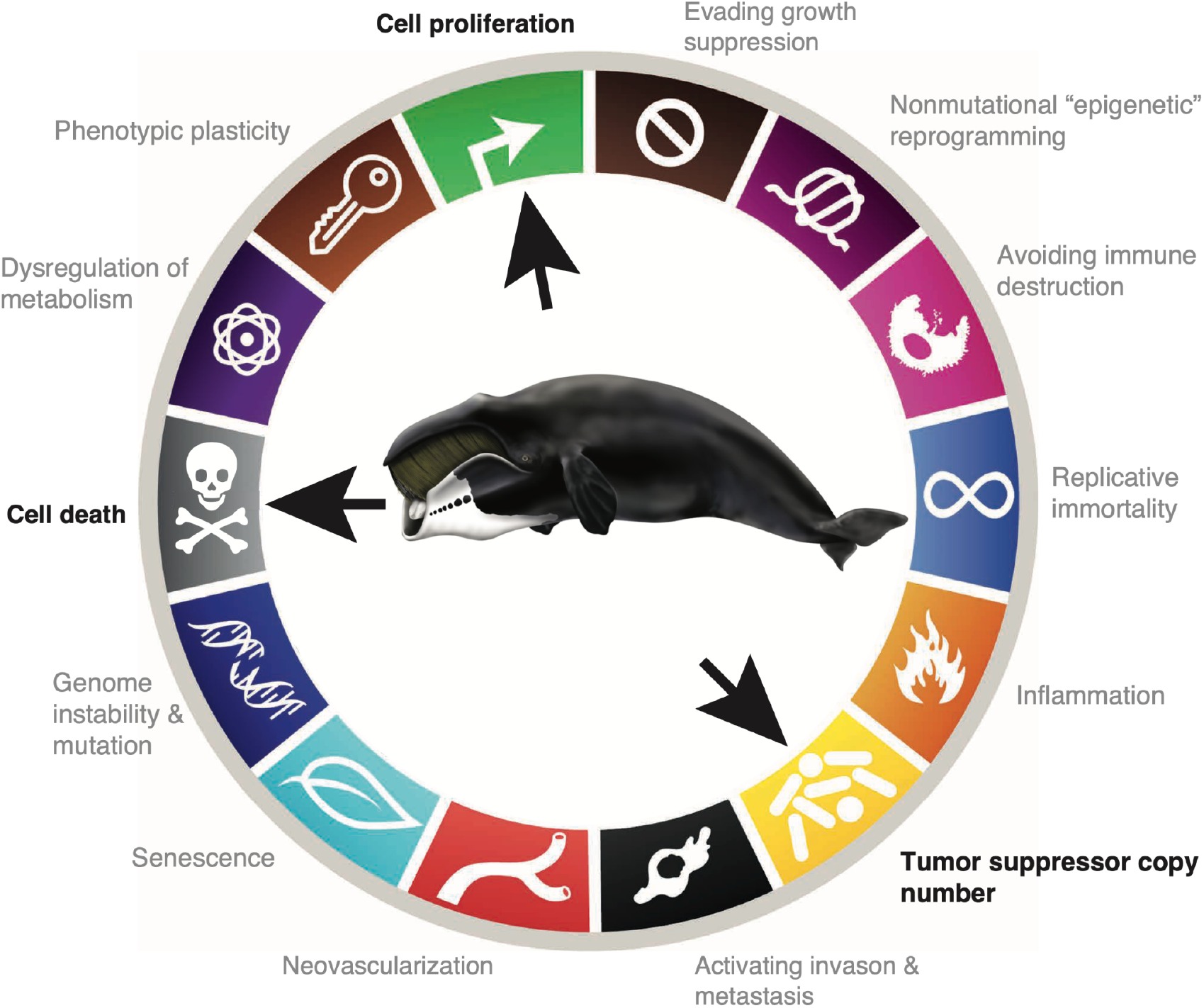

## Introduction

A major constraint on the evolution of long lifespans is an increased risk of developing cancer because organisms with long lifespans have more time to accumulate cancer-causing mutations than organisms with shorter lifespans (Armitage and Doll, 1954; Caulin et al., 2015; Caulin and Maley, 2011; Doll, 1971; Peto, 1975, 2015). The maximum lifespan among vertebrates, for example, ranges from over 400 years in the Greenland shark (Nielsen et al., 2016) to only 59 days in the pygmy goby (Depczynski and Bellwood, 2005). Thus, Greenland sharks have more time to acquire cancer causing mutations than pygmy gobies and should therefore have a higher risk of developing cancer than pygmy gobies. Indeed there is a strong correlation between lifespan and cancer risk within species, for example, cancer prevalence increases with age in humans and rodents (Anisimov et al., 2005) and human genetic variants that predispose to cancer are associated with decreased lifespan (Liu et al., 2022). Conversely, offspring of longer lived parents have a substantially lower incidence of cancer, as well as other common diseases, than offspring of shorter lived parents (Dutta et al., 2013). There is no correlation, however, between maximum lifespan and cancer risk across species (Abegglen et al., 2015; Boddy et al., 2020; Bulls et al., 2022; Vincze et al., 2022) – this lack of correlation is often referred to as ‘Peto’s Paradox’ (Peto, 1975, 2015).

Among the mechanisms long lived animals may have evolved that resolve Peto’s paradox are a decrease in the copy number or functional activity of oncogenes, an increase in the copy number or functional activity of tumor suppressor genes (Caulin et al., 2015; Leroi et al., 2003; Sulak et al., 2016a; Vazquez et al., 2018; Vazquez and Lynch, 2021), the origin of novel tumor suppressors, or increased checks on cell growth, among many others (Gorbunova et al., 2012; Katzourakis et al., 2014; Nagy et al., 2007; Tian et al., 2019, 2013). Mice with extra copies of the master tumor suppressor *TP53* (’super p53’ mice) or the cell cycle checkpoint regulator *CDKN2A* (‘super Ink4a/Arf’ mice), for example, are cancer resistant and age normally, whereas mice with extra copies of *TP53* and *CDKN2A* are cancer resistant and have extended lifespans (Garcia-Cao et al., 2002; Matheu et al., 2007, 2004). Aging is also suppressed in mice that overexpress the tumor suppressor *Klotho*, a peptide hormone that antagonizes the IGF-1 signaling pathway (Kurosu et al., 2005; Xie et al., 2013). Similarly while transgenic mice with an increased dosage of *SIRT1*, a protein deacetylase that contributes to telomere maintenance (Palacios et al., 2010), have normal lifespans they are protected from the development of aging-associated diseases such as glucose intolerance, osteoporosis and cancer (Herranz et al., 2010). These data indicate that mutations which increase copy number or functional dosage can inhibit tumor growth, promote healthy aging, and increase lifespan (Pinkston et al., 2006).

In this study we use evolutionary genomics and comparative experimental biology to explore the mechanisms by which Bowhead whales (*Balaena mysticetus*), which can live which live at least 211 years unlike all other Cetaceans which live 50-70 years (George et al., 2011, 1999; O’Hara et al., 2002; Tacutu et al., 2013), evolved their extremely long lifespans. We found that the Bowhead whale genome encodes a recently duplicated *CDKN2C* (p18/INK4C) gene, unlike all other Sarcopterygians including their closest living relative the Right whale (*Eubalaena sp*.). The duplicate *CDKN2C* is a retrogene (*CDKN2CRTG*) embedded within a Cetacean-specific LINE L1 element, and is highly expressed in Bowhead whale tissues likely because it coopted a L1 promoter to drive constitutive expression. Furthermore we use a series of gain of function experiments to show how the duplicate *CDKN2CRTG* gene may influence cellular phenotypes such as the cell cycle and DNA damage repair in ways that are beneficial for aging and cancer resistance. Remarkably, Bowhead and Right whales only diverged ~4-5 million years ago (Árnason et al., 2018; Dornburg et al., 2012; McGowen et al., 2009), indicating the long lifespan of Bowheads evolved relatively recently and coincident with the origin of *CDKN2CRTG*.

## Results

### Extremely low cancer prevalence in Cetaceans

Sporadic cases of cancer have been reported in many Cetaceans, including several dolphin species, sperm, sei, humpback, beluga, and fin wales, and even remarkably large blue whales and long-lived bowhead whales, (Geraci et al., 1987; Uys and Best, 1966)(Migaki and Albert, 1982; Stimmelmayr et al., 2017)(Geraci et al., 1987; Uys and Best, 1966). Unfortunately, however, while several studies have recently reported cancer prevalence data from diverse mammals these studies were based mostly on zoo animals and therefore did not include Cetaceans. To explore cancer prevalence in Cetacea, we gathered data from the literature and found several published studies with multiple necropsy reports for individual species as well as studies that combined data from multiple species (**Figure 1A**; **Figure 1 – source data 1**). Unexpectedly, among these were two reports that described neoplasias from 7/211 (Migaki and Albert, 1982; Stimmelmayr et al., 2017)) and 18/382 (R. et al., 2020) subsistence-harvested Bowhead whales conservatively suggesting cancer prevalence in Bowheads to be 3.3% (95% CI: 1.3–6.7%) and 4.7% (95% CI: 2.8–7.3%), respectively; By far the most common tumor reported was hepatic lipoma (84%). Other studies, which had very large samples but that combined data from multiple species, reported cancer in only 14/1800 (0.78%, 95% CI: 0.04-0.13%) and 2/2000 (0.01%, 95% CI: 0.00–0.04%%) necropsies (**Figure 1A**; **Figure 1 – source data 1**). Next, we compared neoplasia prevalence in Cetacea to other Therian mammals from three published studies that included pathology reports from 37 (Boddy et al., 2020) and 191 (Vincze et al., 2022) species of Therian mammals, and a large dataset of Xenarthrans compiled from multiple published sources (Vazquez et al., 2022); The dataset includes cancer prevalence data from 244 species. As a group the mean prevalence of neoplasia in Cetacea was 1.4% (95% CI: 1.07– 1.70%) when aggregated across all datasets and excluding St. Lawrence estuary belugas (**Figure 1B**; **Figure 1 – source data 2**). By rank, Cetaceans are the second most cancer resistant mammalian order (**Figure 1B**; **Figure 1 – source data 2**). Thus, cancer occurs but is also likely extremely uncommon in Cetaceans including Bowhead whales.

**Figure 1.**
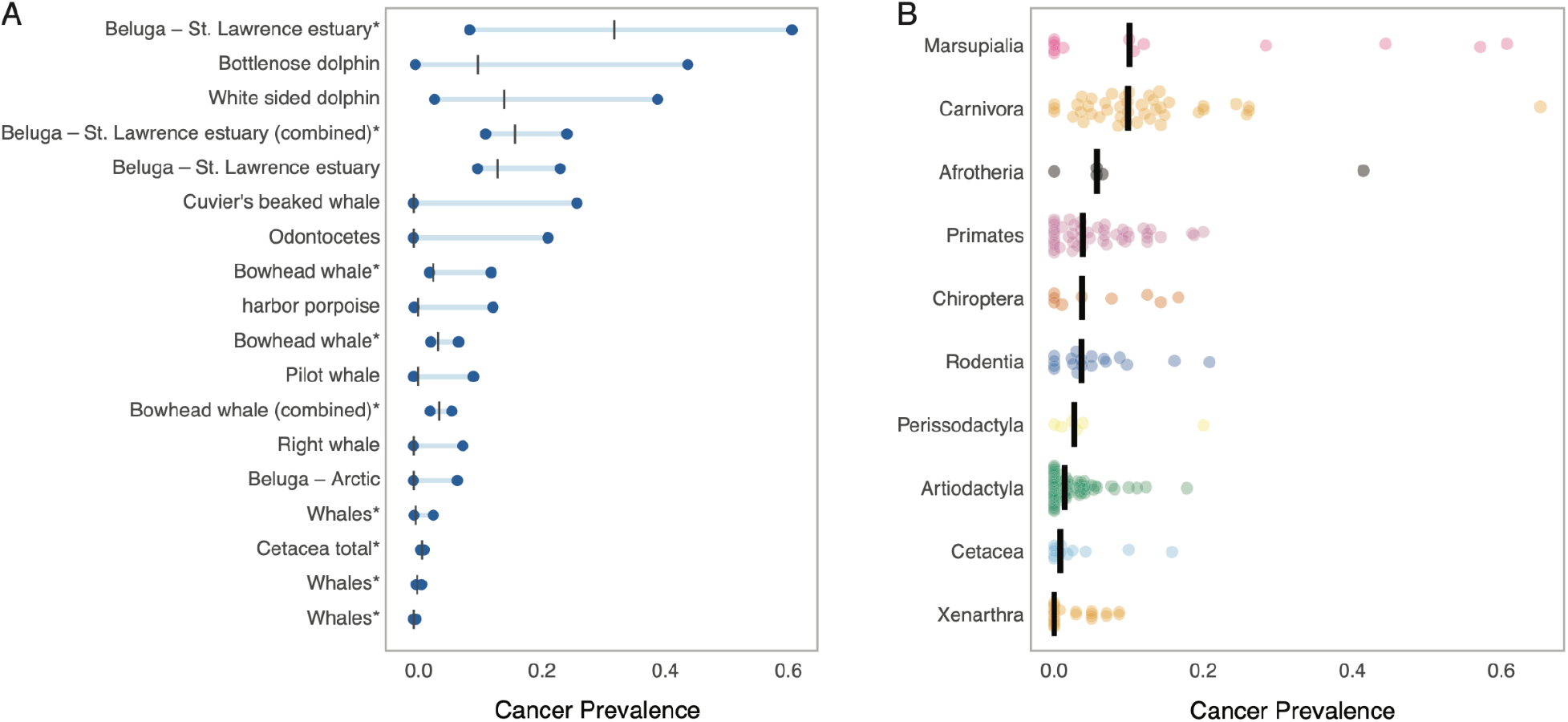
Extremely low cancer prevalence in Cetaceans. **(A)** Lolliplot plot showing the mean (black bar) and Clopper-Pearson exact 95% confidence intervals of cancer prevalence in each group, asterisks (*) indicate taxa with more than 82 pathology reports. Data from more than two studies are indicated with “(combined)”. **(B)** Cancer prevalence data for each species (dot) grouped by clade and shown as jittered stripchart. The number of species per clade is: Afrotheria=5, Artiodactyla=68, Carnivora=43, Cetacea=4, Chiroptera=9, Marsupialia=13, Perissodactyla=6, Primates=47, Rodentia=18, and Xenarthra=14. Medians for each clade are shown with vertical black bars. **Figure 1 – source data 1**. Cancer prevalence shown in panel A. **Figure 1 – source data 2**. Cancer prevalence shown in panel B.

### The Bowhead whale genome encodes a duplicate *CDKN2C* retrogene (*CDKN2CRTG*)

Whales, particularly long-lived Bowheads, must have evolved cancer protection mechanisms that reduce the enhanced risk of cancer imposed by their large bodies and long lifespans. Previous studies, for example, identified several gene duplications in the Bowhead whale genome associated with cancer biology (Keane et al., 2015; Tejada-Martinez et al., 2021). These studies, however, used methods that may not be able to identify gene duplications that have very low sequence divergence and did not include Right whales as an outgroup. Thus, they were unable to identify very recent duplications, such as those that occurred since the Bowhead-Right whale divergence ~4-5 MYA, or Bowhead whale specific duplications. To identify duplicates with low sequence divergence, we used reciprocal best BLAT (RBB) and refined our copy number estimates based on total copy coverage (ECNC, (Vazquez and Lynch, 2021)) to and duplications fragmented in lower quality genomes (Vazquez and Lynch, 2021) to identified gene duplications in the genome of a Bowhead whale, a South Atlantic Right whale, a North Atlantic Right whale, and a sperm whale (**Figure 2A**; **Figure 2 – source data 1**). Intersection of these duplicate gene sets identified a single duplication unique to the Bowhead whale genome: *Cyclin Dependent Kinase Inhibitor 2C* (*CDKN2C*; **Figure 2B**), which encodes a member of the INK4 family of cyclin-dependent kinase inhibitors that regulate the G_1_/S checkpoint of the cell cycle (Cui et al., 2015).

**Figure 2.**
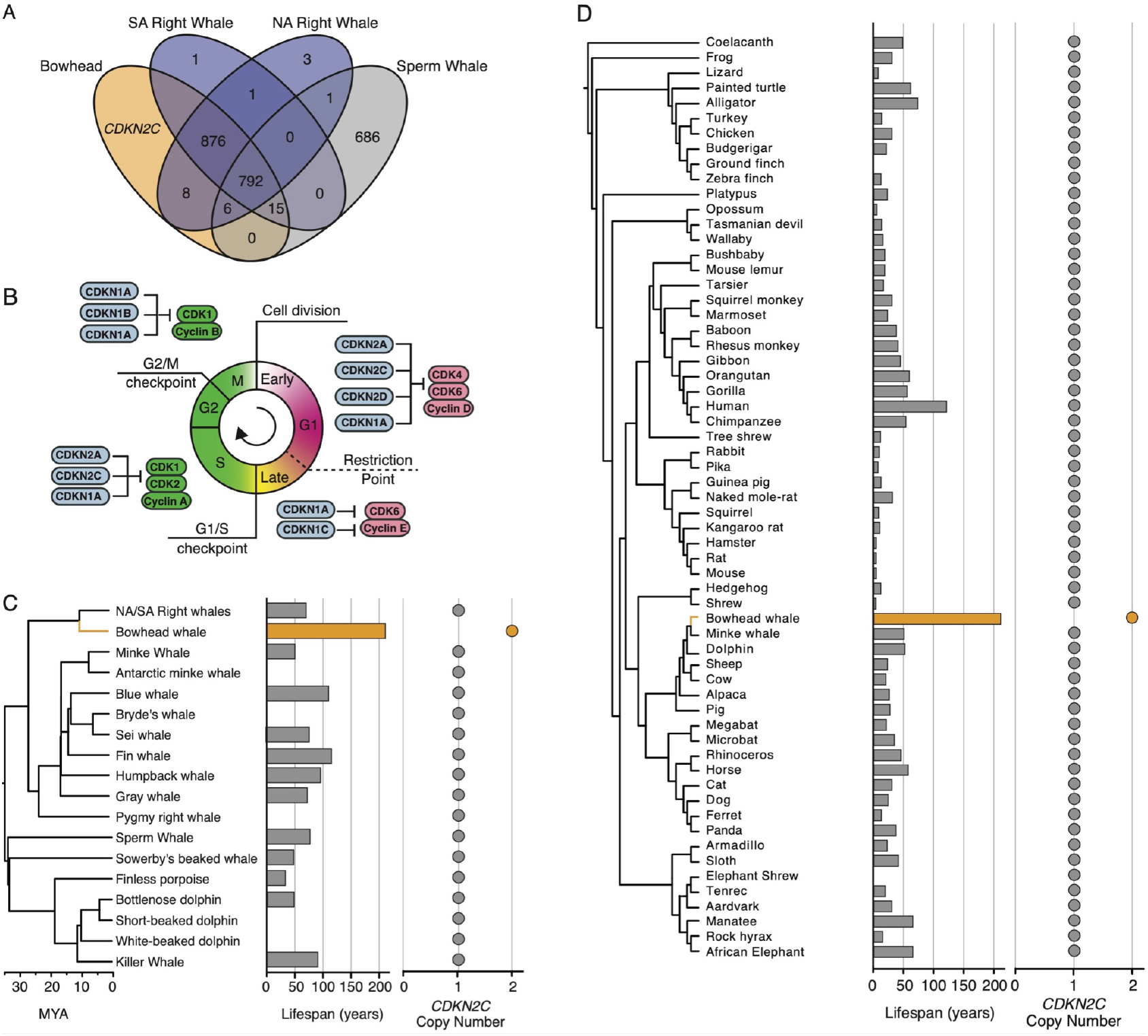
Duplication of the cell-cycle regulator *CDKN2C* in Bowhead whales. **(A)** Venn diagram showing intersection of gene duplicated between Bowhead whale, South Atlantic (SA) Right whale, North Atlantic (NA) Right whale, and Sperm whale. Note that only 1 gene duplicate is unique to Bowhead whales: *CDKN2C*. **(B)** Cartoon of the cell cycle, with major regulatory genes for each phase. **(C)** Lifespan and *CDKN2C* copy number in 18 Cetaceans. Cetacean phylogenetic tree is scaled to time. Lifespan data is from (de Magalhães and Costa, 2009). **(D)** Lifespan and *CDKN2C* copy number in 61 Sarcopterygians. Lifespan data is from (de Magalhães and Costa, 2009). **Figure 2 – source data 1**. Duplicate genes identified in the Bowhead whale, North and South Atlantic Right whales, and Sperm whale. **Figure 2 – source data 2**. Sarcopterygian genomes characterized for *CDKN2C* copy number.

To verify that the *CDKN2C* gene is uniquely duplicated in Bowhead whales, we manually verified *CDKN2C* copy number in the genomes of 80 Sarcopterygians including 19 Cetaceans. We found that all Cetacean (**Figure 2C**) and Sarcopterygian genomes (**Figure 2D**; **Figure 2 – source data 2**) encoded a single *CDKN2C* gene except the Bowhead whale, which we confirmed encoded a typical *CDKN2C* gene as well as a *CDKN2C* retrogene (*CDKN2CRTG*), i.e. an intronless *CDKN2C* gene, and the allotetraploid African clawed frog (*Xenopus laevis*) which has previously been reported to have two *CDKN2C* genes (CDKn2c.L and CDKn2c.S) (Tanaka 2017). *CDKN2C* encodes two transcripts that initiate from alternative promoters (**Figure 3A**). While both transcripts contain coding exons 2 and 3, transcripts initiated from the distal promoter contain an alternative noncoding exon (exon 1) that results in ~1-2kb 5’-UTR whereas transcripts generated from the proximal promoter have a ~400bp 5’-UTR (Blais et al., 1998; Phelps et al., 1998). The long transcript is predominantly expressed in germ cells (Buchold et al., 2007; Franklin and Xiong, 1996), suggesting the long transcript is more likely to be retroduplicated than the short one. Indeed, the *CDKN2CRTG* gene includes non-coding exon 1, indicating it was reverse transcribed from the long *CDKN2C* transcript (**Figure 3B**). As expected for a recent LINE L1 mediated retrotransposition event, the *CDKN2CRTG* gene includes remnants of a long poly-A tail, and is flanked by a LINE1 target site (GT/AAAA) and direct repeats (**Figure 3B**).

**Figure 3.**
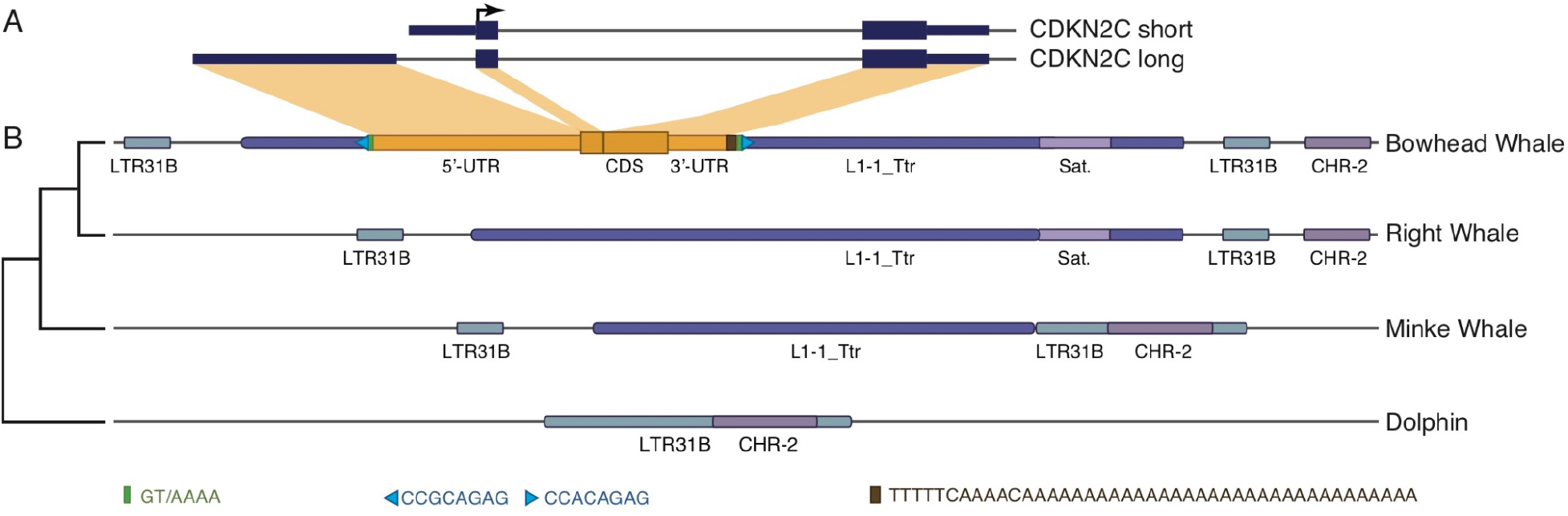
The Bowhead whale duplicate *CDKN2C* is a retrogene (*CDKN2CRTG*) embedded within a LINE L1 element. **(A)** The canonical *CDK2NC* gene encodes two isoforms that use alternative 3’-UTR exons. **(B)** Structure of the locus encoding *CDK2NCRTG* gene in Bowhead whale, and the homologous locus in Right whale, Minke whale, and Dolphin (phylogeny shown on left). The *CDK2NCRTG* gene is embedded within a Cetacean-specific LINE L1 element (L-1_Ttr) and nearby other transposable elements. The location of the poly-A tail (brown rectangle), LINE1 target site (green rectangle), and direct repeats (blue triangle) are shown. The Regions of the *CDK2NC* long transcript that contribute to the *CDK2NCRTG* gene are highlighted in yellow.

The *CDKN2CRTG* gene is embedded within the 5’-UTR of a Cetacean-specific LINE L1 (L1-2_Ttr) that is flanked by a unique combination of transposable elements that allow us to annotate the locus in other Cetacean genomes (**Figure 3B**). The orthologous locus in other whale genomes, including North and South Atlantic Right whales, Minke whale, and Dolphin do not contain *CDKN2CRTG* (**Figure 3B**). The Bowhead whale *CDKN2C* and *CDKN2CRTG* genes are 100% identical across the 5’-UTR, coding region, and 3’-UTR. These data indicate that the retroduplication event occurred very recently in the Bowhead-lineage, coincident with the evolution of a long lifespan, and may have generated a functional *CDKN2C* gene.

### *CDKN2CRTG* is transcribed in multiple tissues

If retoduplication of the *CDKN2C* gene played a role in the evolution of long lifespans in Bowhead whales, then *CDKN2CRTG* should be transcribed. Therefore, we used previously published RNA-Seq data from seven Bowhead whale tissues (Keane et al., 2015), five Minke whale tissues, (Yim et al., 2014), one Humpback whale tissue, and one Sperm whale tissue, and 454 GS FLX data from five dolphin tissues (Westbury et al., 2019) to determine if *CDKN2CRTG* was transcribed and compared with *CDKN2C* expression levels (**Figure 4 – source data 1**). We found the *CDKN2CRTG* transcripts from Bowhead tissues spanned the junction between the *CDKN2CRTG* 3’-end and the L1-2_Ttr LINE element (**Figure 4A**), allowing us to discriminate between *CDKN2C* and *CDKN2CRTG* transcripts. *CDKN2CRTG* transcripts were on average ~27x more abundant than *CDKN2C* transcripts (**Figure 4B**), which were expressed at relatively low levels or not at all (**Figure 4B**). We note that cDNA libraries from Bowhead tissues were constructed from poly-A captured RNA, indicating the *CDKN2CRTG* transcript is likely polyadenylated.

**Figure 4.**
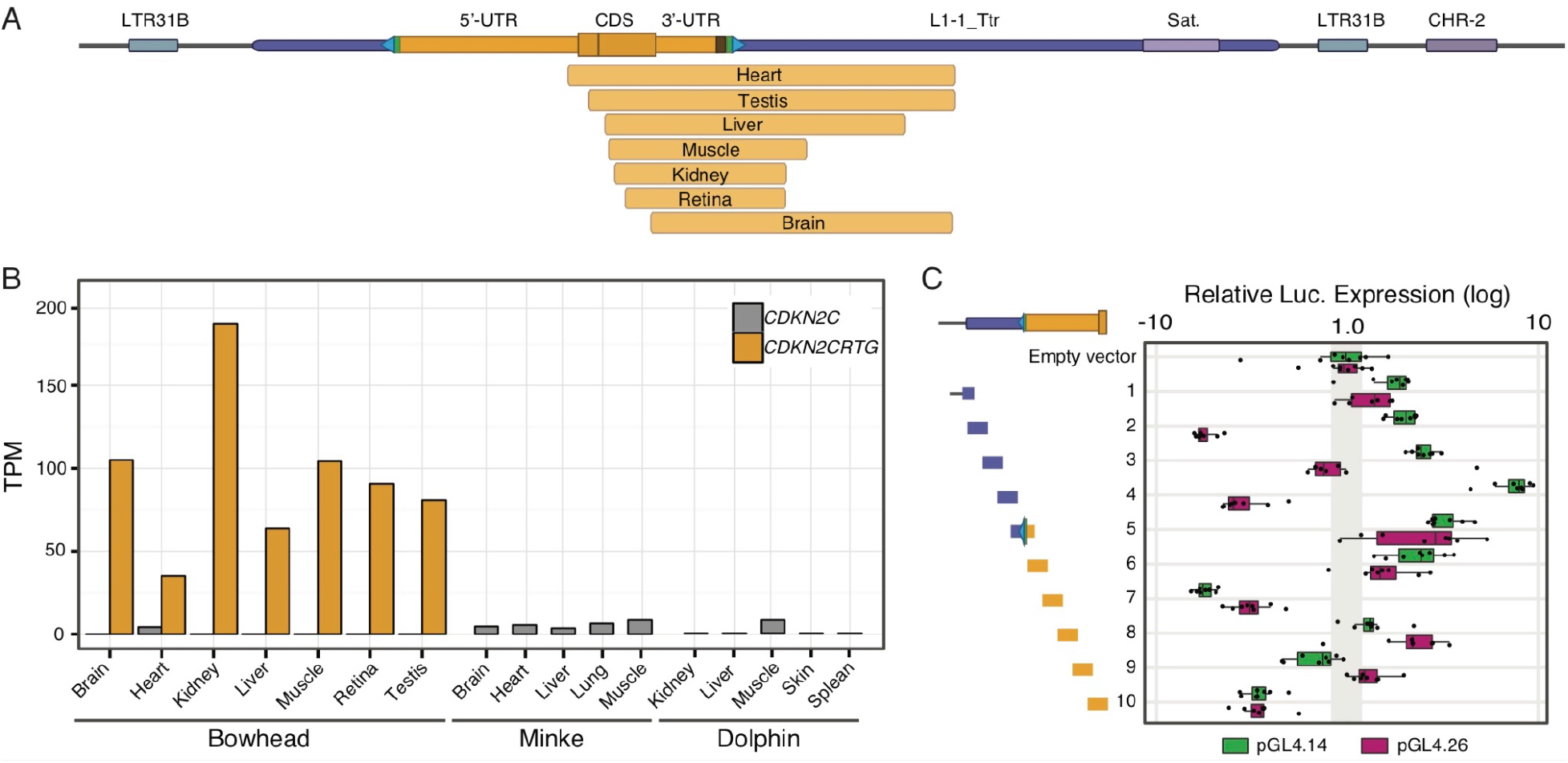
*CDKN2CRTG* **is expressed and from a coopted LINE L1 promoter**. **(A)** Upper, *CDKN2CRTG* locus. Lower, *CDKN2CRTG* transcripts reconstructed with HISAT2 and StringeTie. Note that all transcripts extend downstream into the LINE L1 L2-1_Ttr and include the junction between the *CDKN2CRTG* gene and the ctranposable element which allows for discrimination between *CDKN2CRTG* and *CDKN2C* transcripts. **(B)** Expression of *CDKN2CRTG* (yellow) and *CDKN2C* (grey) in Bowhead whale, Minke whale, and Dolphin tissues. Transcript abundances in transcripts per million (TPM) were inferred with Kallisto. **(C)** Dual luciferase reporter assay in transiently transfected mouse 3T3 cells from ten 301bp overlapping fragments that tile the region from upstream of the L1-2_Ttr element to downstream of the *CDKN2CRTG* initiation codon. Each fragment was cloned into luciferase reporter vector pGL4.14[*luc2*/Hygro], which lacks both an enhancer and a promoter, and pGL4.26[*luc2*/minP/Hygro], which lacks an enhancer but has a minimal promoter. Relative luciferase expression (standardized to *Renilla* and empty vector control), n=8. **Figure 4 – source data 1. Transcriptome sample information.** **Figure 4 – source data 2. Data shown in panel C.**

### A Coopted LINE L1 promoter drives *CDKN2CRTG* transcription

Most retrogenes lack native regulatory elements such as promoters and enhancers that direct their expression, thus expression of *CDKN2CRTG* is either regulated by existing regulatory elements or ones that evolved *de novo*. Our observation that the *CDKN2CRTG* gene is embedded within a LINE L1 element suggests that *CDKN2CRTG* transcription may be regulated by the endogenous LINE L1 promoter/enhancer located within the LINE L1 5’-UTR (Lavie et al., 2004; Swergold, 1990). To test if this region has regulatory abilities, we synthesized ten 301bp fragments including 51bp 5’- and 3’-overlaps that tile the region from upstream of the L1-2_Ttr element to downstream of the *CDKN2CRTG* initiation codon (**Figure 4C**) and cloned each fragment into the pGL4.14[*luc2*/Hygro] and pGL4.26[*luc2*/minP/Hygro] luciferase reporter vectors; the pGL4.14[*luc2*/Hygro] vector lacks an enhancer and a promoter whereas the pGL4.26[*luc2*/minP/Hygro] vector contains a minimal promoter allowing us to test the cloned fragments for promoter and enhancer functions. Next, we transiently transfected Chinese Hamster Ovary (CHO) cells with each reporter construct and used a dual luciferase reporter assay to quantify luciferase expression. Fragments from the 5’-UTR of the L1-2_Ttr element significantly increased luciferase expression from the pGL4.14[*luc2*/minP/Hygro] reporter vector peaking with fragment four, which increased luciferase expression 87-fold (95.0%CI 70.7-107 fold) (**Figure 4C** and **Figure 4 – figure supplement 1**). In contrast, these fragments did not increase luciferase expression when cloned into the pGL4.26[*luc2*/minP/Hygro] vector (**Figure 4C** and **Figure 4 – figure supplement 2**), suggesting this region can function as a promoter but likely not an enhancer. Fragments five and six, however, which include the target site and the most proximal region of the *CDKN2CRTG* 5’-UTR, did increase luciferase expression from the pGL4.26[*luc2*/minP/Hygro] vector (**Figure 3C** and **Figure 3 – figure supplement 2**) indicating this region has enhancer/promoter potential. These data suggest that expression of *CDKN2CRTG* is regulated by an enhancer/promoter located within the LINE L1-2_Ttr element immediately upstream *CDKN2CRTG* 5’-UTR.

### *CDKN2CRTG* overexpression induces cell-cycle delay at the G_1_ and G_1_/S checkpoints

*CDKN2C* expression is cell-cycle dependent with transcription initiating around the late G_1_/S checkpoint, peaking in the S and G_2_/M phases before falling again in early G_1_ (Hirai et al., 1995) and acts as a G_1_/S checkpoint (Cui et al., 2015). However, the high level of *CDKN2CRTG* expression and the usage of LINE L1 derived promoter to drive expression suggests constitutive expression of *CDKN2CRTG* throughout the cell-cycle. To explore the consequences of constitutively high *CDKN2CRTG* expression on cell cycle dynamics, we first generated a stable Fucci-2a reporter CHO cell line (FuCHO). The Fucci-2a system consists of a monomeric Cherry (mCherry) tagged fragment of human CDT1 (hCdt1) gene and a monomeric Venus (mVenus) tagged fragment of human GMNN (hGem) gene joined by the *Thosea asigna* virus 2A (T2A) self-cleaving peptide (Mort et al., 2014; Sakaue-Sawano et al., 2008) (**Figure 5A**). After transcription, the T2A peptide cleaves generating an mCherry-hCdt1(30/120) probe that accumulates during G_1_ phase and is degraded at the G_1_-S transition and a mVenus-hGem probe that accumulates during S/G_2_/M phases and is rapidly degraded prior to cytokinesis (**Figure 5B**). Next we synthesized and cloned the *CDKN2CRTG* gene into the mammalian pcDNA3.1-C-eGFP expression vector, which adds a C-terminal eGFP tag, transiently transfected this vector into FuCHO cells, and quantified the cell-cycle stage using flow cytometry. Consistent with previous studies of CDKN2C, we found that overexpression of *CDKN2CRTG* induced an accumulation of cells in G_1_ (Hirai et al., 1995) and at the G_1_/S checkpoint (Cui et al., 2015), which was particularly pronounced in the GFP+ population of FuCHO cells (**Figure 5C-E**). These results indicate that the bowhead whale CDKN2CRTG protein is functionally equivalent to CDKN2C with respect to its ability to negatively regulate the cell cycle during the G_1_ phase. .

**Figure 5.**
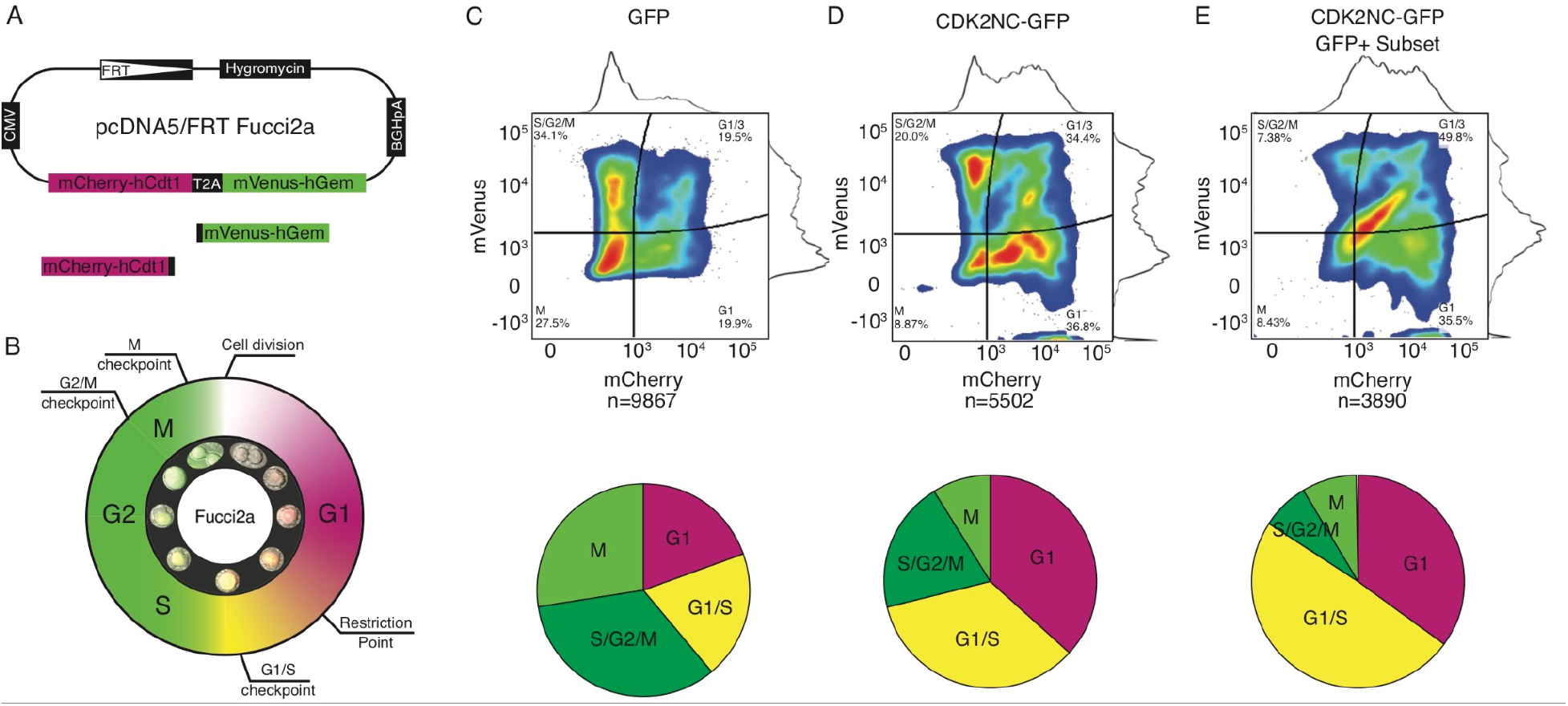
CDK2CRTG overexpression induces cell cycle delay at G_1_ and G_1_/S. **(A)** Fucci2a Flp-In reporter used to generate the FuCHO cell line. **(B)** Schematic of the cell cycle as visualized with the mCherry-hCdt1 and mVenus-hGem fluorescent reporters. Inset shows fluorescence intensity of single FuCHO cell. Cell cycle stage in FuCHO cells either mock transfected or transfected with a GFP-tagged copy of the Bowhead *CDKN2CRTG* was assayed after 20 hours by flow cytometry. **(C)** The distribution of cell cycle states in mock transfected FuCHO cells (upper; inset percentages indicate the frequency of cells in each stage), pie chart (lower) shows the percentage of cells in each cell cycle stage. **(D)** The distribution of cell cycle states in *CDKN2CRTG* transfected FuCHO cells (upper; inset percentages indicate the frequency of cells in each stage), pie chart (lower) shows the percentage of cells in each cell cycle stage. Note the shift in cells towards G1 and G1/S phase compared to mock transfected cells (shown in panel C). **(E)** The distribution of cell cycle states GFP gated *CDKN2CRTG* transfected FuCHO cells. Note the shift in cells towards G1 and G1/S phase compared to mock transfected cells (shown in panel C). The shift in the population to G1 and G1/S is particularly pronounced compared to D.

### *CDKN2CRTG* overexpression protects cells from DNA damage induced apoptosis

Ectopic overexpression of *CDKN2C* can induce cell cell arrest and apoptosis in normal and cancerous mouse and human cells (Komata et al., 2003a; Kulkarni et al., 2002; Schreiber et al., 1999), suggesting that a duplicate *CDKN2C* gene expressed at high levels may also affect DNA damage processes. Therefore, we used an ApoTox-Glo assay to explore the effects of CDKN2CRTG-eGFP overexpression in FuCHO cells treated with doxorubicin for 18 hours. Doxorubicin, a topoisomerase poison, induces DNA double stranded breaks by intercalating with DNA and inhibiting the progression of topoisomerase II complex during transcription (Pommier et al., 2010) and during DNA replication in S phase (Tacar et al., 2013). As expected, doxorubicin treatment alone induced apoptosis (**Figure 6A**) and decreased cell viability (**Figure 6B**) whereas CDKN2CRTG overexpression protected cells from apoptosis and dramatically preserved cell viability (**Figure 6A/B**). Thus, CDKN2CRTG expression can prevent the accumulation of DNA damage.

**Figure 6.**
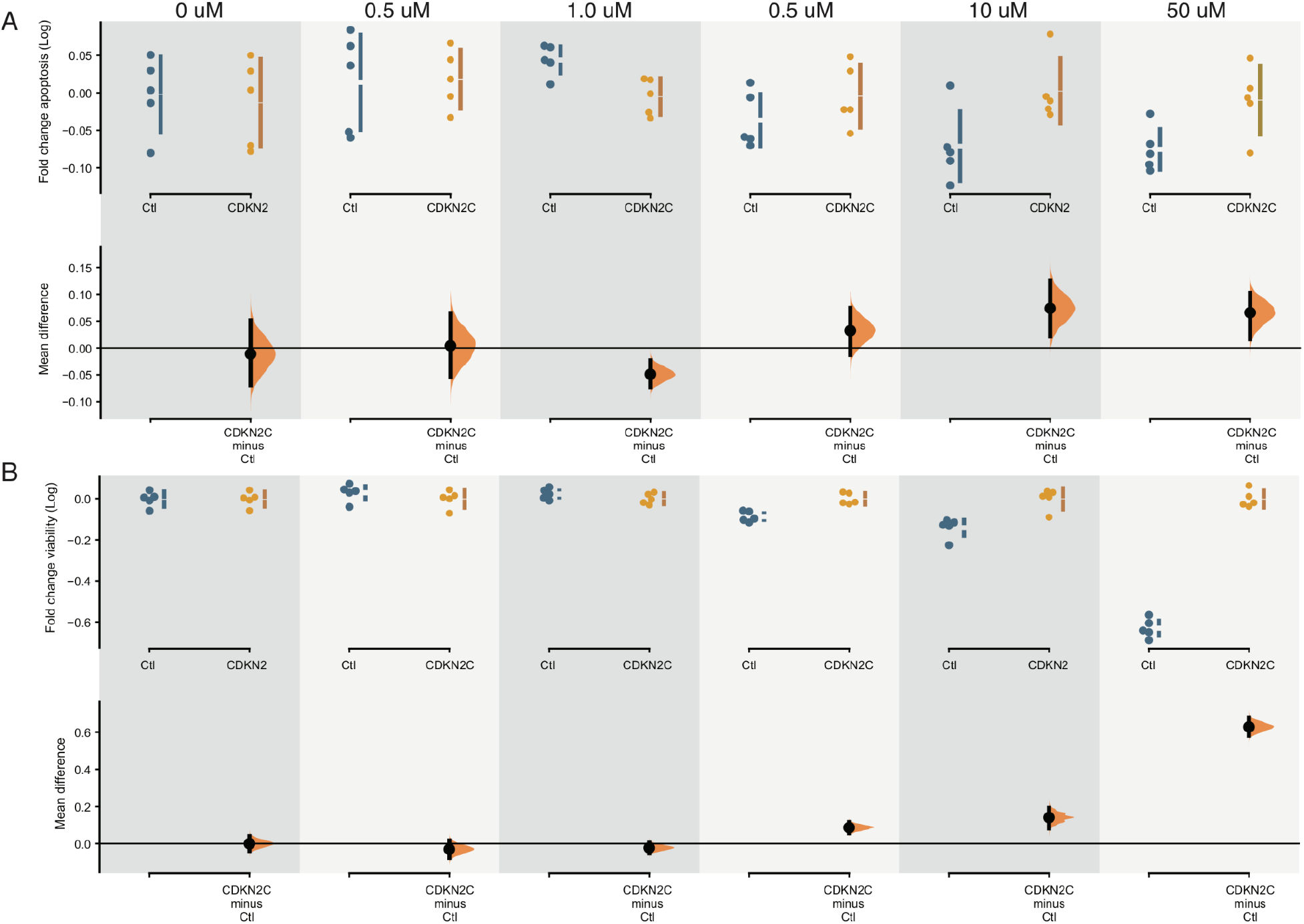
CDK2CRTG overexpression protects FuCHO cells from doxorubicin induced apoptosis and cell death. ApoTox-Glo results for control (Ctl) and CDKN2C transfected FuCHO cells treated with increasing doses of doxorubicin (0-50 uM) for 18 hours. Fold change in apoptosis (A) and viability (B) are shown on the upper axes of each plot. On the lower axes, mean differences are plotted as bootstrap sampling distributions. Each mean difference is depicted as a dot. Each 95% confidence interval (CI) is indicated by the ends of the vertical error bars. The effect sizes and CIs are reported above as: effect size [CI width lower bound; upper bound]. 5000 bootstrap samples were taken; the confidence interval is bias-corrected and accelerated. The P-value(s) reported are the likelihood(s) of observing the effect size(s), if the null hypothesis of zero difference is true from a two-sided permutation t-test. **(A)** Apoptosis results: The unpaired mean difference between Ctl-0 and CDKN2C-0 is −0.0111 [95.0%CI −0.0695, 0.051], P=0.8. The unpaired mean difference between Ctl-0.5 and CDKN2C-0.5 is 0.00405 [95.0%CI −0.0535, 0.0643], P=0.878. The unpaired mean difference between Ctl-1 and CDKN2C-1 is −0.0488 [95.0%CI −0.0727, −0.0231], P=0.0218. The unpaired mean difference between Ctl-5 and CDKN2C-5 is 0.0322 [95.0%CI −0.013, 0.0746], P=0.24. The unpaired mean difference between Ctl-10 and CDKN2C-10 is 0.0738 [95.0%CI 0.0222, 0.125], P=0.0292. The unpaired mean difference between Ctl-50 and CDKN2C-50 is 0.0657 [95.0%CI 0.0164, 0.102], P=0.0188. **(B)** Cell viability results: The unpaired mean difference between Ctl-0 and CDKN2C-0 is −0.000142 [95.0%CI −0.0401, 0.04], P=0.984. The unpaired mean difference between Ctl-0.5 and CDKN2C-0.5 is −0.0301 [95.0%CI −0.0769, 0.0166], P=0.294. The unpaired mean difference between Ctl-1 and CDKN2C-1 is −0.0227 [95.0%CI −0.0518, 0.00616], P=0.192. The unpaired mean difference between Ctl-5 and CDKN2C-5 is 0.0863 [95.0%CI 0.0569, 0.116], P=0.0014. The unpaired mean difference between Ctl-10 and CDKN2C-10 is 0.141 [95.0%CI 0.0823, 0.192], P=0.0002. The unpaired mean difference between Ctl-50 and CDKN2C-50 is 0.628 [95.0%CI 0.581, 0.679], P=0.0. **Figure 6 – source data 1**. Data shown in panels A and B.

### Bowhead whale cells have longer doubling times than other whale cells

The role of CDKN2C in regulating the G_1_ and G_1_/S cell cycle checkpoints suggest that Bowhead whale cells may have different cell cycle dynamics than other species. While measurements of cell cycle phase length or overall duration are not available for most species, many studies report population doubling times from mammalian cells in culture. Population doubling time is proportional to the cumulative length of individual cell cycle phases, particularly the length of G_1_ (Blank et al., 2018). Therefore, we performed a thorough literature review and assembled a dataset of population doubling times from 60 species of Eutherian mammals (Vazquez et al., 2022). While there was considerable variation in population doubling times between species, whale cells had similar doubling times as other species of mammals (**Figure 7A**). Bowhead whale cell lines (n=4), however, had significantly longer doubling times than cell lines from other whales (n=13) with an unpaired mean difference in doubling time of 15.4 hours (95.0%CI 9.46-24.9) and a two-sided permutation t-test P-value=0.0104 (**Figure 7B**). Bowhead whale cell lines also had longer doubling times than cell lines from Right whales (n=3), with an unpaired mean difference in doubling time of 14.2 hours (95.0%CI 7.58-25.9) and a two-sided permutation t-test P-value=0.0 (**Figure 7C**). We note that these differences in population doubling times may reflect biological differences between species or technical variation not related to species-specific differences in doubling time or cell cycle length.

**Figure 7.**
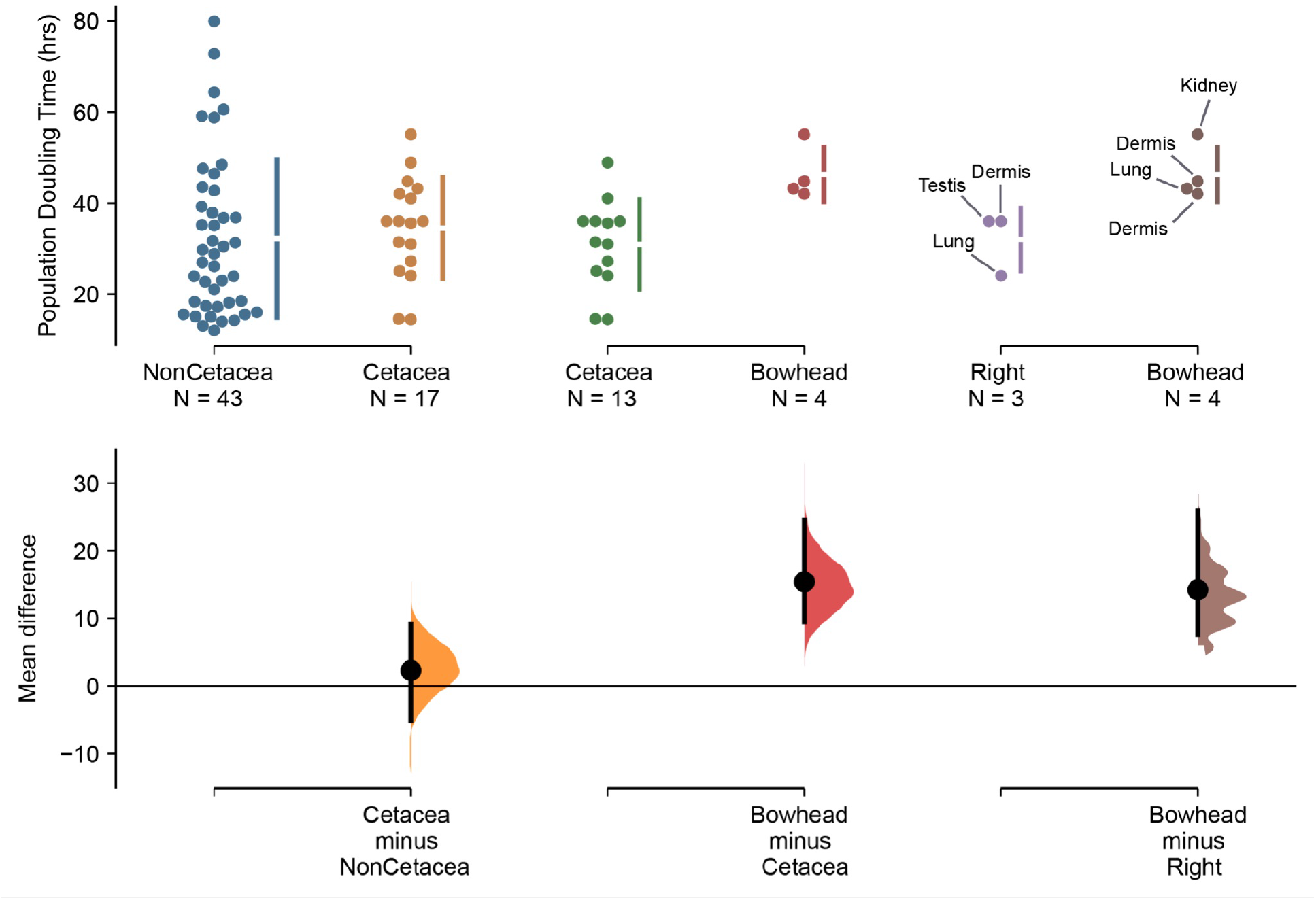
Bowhead whale cells have longer doubling times than cells from other whales. Population doubling times from 60 species of Eutherian mammals collected from the literature. Population doubling times for each group is shown on the upper axis (along with cell-types from which doubling times were estimated for Right and Bowhead whales). On the lower axes, mean differences are plotted as bootstrap sampling distributions. Each mean difference is depicted as a dot. Each 95% confidence interval (CI) is indicated by the ends of the vertical error bars. The effect sizes and CIs are reported above as: effect size [CI width lower bound; upper bound]. 5000 bootstrap samples were taken; the confidence interval is bias-corrected and accelerated. The P-value(s) reported are the likelihood(s) of observing the effect size(s), if the null hypothesis of zero difference is true from a two-sided permutation t-test. The unpaired mean difference between NonCetacea and Cetacea is 2.28 [95.0%CI −5.12, 9.14], P=0.616. The unpaired mean difference between Cetacea and Bowhead is 15.4 [95.0%CI 9.46, 24.6], P=0.0104. The unpaired mean difference between Right and Bowhead is 14.2 [95.0%CI 7.58, 25.9], P=0.0. Cell-types are shown for Bowhead and Right Whale cell lines. **Figure 7 – source data 1. Data shown in Figure 7**.

## Discussion

Among the challenges for discovering the genetic and molecular changes associated with the evolution of remarkably long lifespans is identifying long-lived species that are closely related to species with ‘normal’ body sizes and lifespans. The ideal study system for exploring how animals evolved long healthy lifespans, for example, would be one in which an exceptionally long-lived lineage was deeply nested within a clade with short lifespans, for which genomes and potentially other biological materials are available for study. The long-lived species should also be very closely related to short-lived species, allowing us to identify derived genetic changes that are phylogenetically associated with lifespan extension rather than other traits or neutral genetic changes. Bowhead whales are an excellent system in which to study lifespan evolution because they can live over 200 years, unlike other whales which live ~60 years, and diverged from Right whales only ~4-5 million years ago (Árnason et al., 2018; Dornburg et al., 2012; McGowen et al., 2009). Thus, Bowhead whales evolved their extreme lifespans relatively recently allowing us to identify potentially causal mutations from a sea of neutral ones.

### Extremely low cancer prevalence in Cetaceans

Previous studies have reported sporadic cases of cancer in many Cetacean species, including several dolphin species, sperm, sei, humpback, beluga, fin, blue, right, and bowhead whales, (Geraci et al., 1987; Migaki and Albert, 1982; Stimmelmayr et al., 2017; Uys and Best, 1966), but there have been few systematic studies of cancer prevalence in whales. To estimate cancer prevalence in Cetaceans, we gathered mortality data from previously published studies and found that in general, the prevalence of cancer among Cetaceans was low – only 1.4% (95% CI: 1.07–1.70%) when aggregated across all datasets and excluding St. Lawrence estuary belugas (**Figure 1**). Remarkably, among these published studies were two that reported neoplasias from 7/211 (Stimmelmayr et al., 2017) and 18/382 (R. et al., 2020) subsistence-harvested Bowhead whales, suggesting that cancer prevalence in Bowheads to be 3.3% (95% CI: 1.3–6.7%) and 4.7% (95% CI: 2.8–7.3%), respectively. Thus, despite their remarkably large sizes, and in the case of Bowhead whales long lifespans, cancer prevalence in Cetaceans is generally very low. Indeed, Cetaceans are the second most cancer resistant mammalian order (**Figure 1**). We note, however, that these estimates are likely to be very dependent on the sample size of necropsy reports. For example, cancer was detected in at least one individual in almost all species with more than 82 individual pathological records illustrating that cancer is likely to be detected in all mammals with adequate sample sizes (Vincze et al., 2022). The number of post-mortem pathology reports was below 82 for most Ceteceans suggesting caution when interpreting the low cancer prevalence in these species, yet these data indicate that whales must have evolved enhanced cancer suppression mechanisms to reduce their cancer risk because in aggregate cancer is relatively rare in Cetaceans.

### What might be the consequences of *CDKN2C* duplication? A longer cell cycle

CDKN2C inhibits cyclin-CDK4/6 complexes (Guan et al., 1994; Noh et al., 1999) that are required for G_0_/G_1_ progression and entry in S phase (Robitaille et al., 2016), thereby acting as a G_1_/S checkpoint (Cui et al., 2015) and regulating the length of G_1_ (Dong et al., 2018). Ectopic overexpression of *CDKN2C*, for example, reduces cell growth by preventing CDK4/6 activation, lengthening G_1_, and delaying progression through the G_1_/S checkpoint (Bench et al., 1996; Komata et al., 2003b; Schreiber et al., 1999; Solomon et al., 2008). In contrast, pharmacological inhibition or knockdown of *CDKN2C* expression promotes cell cycle progression and proliferation (Gao et al., 2015; McClendon et al., 2012). Indeed, we found that overexpression of *CDKN2CRTG* increased the proportion of FuCHO cells in G_1_ and G_1_/S. These data are broadly congruent with the “textbook” model of the eukaryotic cell cycle (Lubischer, 2007), in which the length of the G_1_ phase accounts for most of the variation in cell cycle length between cell-types and species (Blank et al., 2018; Brauer et al., 2008; Fisher, 2016; Johnston et al., 1977; Lubischer, 2007; Wheals, 1985). These data suggest that cell cycle length, particularly the length of the G_1_ phase, may be longer in Bowhead whale cells than cells from other species.

Consistent with this hypothesis, Bowhead whale cells grow extremely slowly (Jarrell, 1979; Smith et al., 1987) prompting us to perform a literature review to compare growth rates of mammalian cells. The most commonly reported measurement for the growth rate of cells in culture is the population doubling time, i.e. the time it takes for a population to double its size (Lindström and Friedman, 2020); while not a direct estimate of cell cycle length, the population doubling time is proportional to cell cycle length and largely determined by the duration of G_1_ (Blank et al., 2018). We found that there was significant variation in doubling time between mammalian cells, and that the mean population doubling time of multiple Bowhead whale cell-types was 14.2 hours (95.0%CI 7.58-25.9) longer than the doubling times of Right whale cells and 15.4 hours (95.0%CI 9.46-24.9) longer than the doubling times of other whale cells. We note that these differences in population doubling times may reflect real biological differences in cell cycle length between species, as well as differences between cell-types, technical differences in how cells were cultured, or how doubling times were calculated, among other coundounfers not directly related to species-specific differences in doubling time or cell cycle length. Yet, these data are consistent with a model in which a consequence of duplicate *CDKN2C* gene, particularly one that is highly expressed, is a decrease in cell growth rates.

### What might be the consequences of *CDKN2C* duplication? Enhanced DNA damage repair

Activation of cell cycle checkpoints in response to DNA damage pauses cell cycle progression, which give cells time to repair damage or induce apoptosis thereby preventing transmission of damaged or incompletely replicated chromosomes (Bartek and Lukas, 2001; O’Connell and Cimprich, 2004; O’Connor, 1997). Previous studies have also found that the INK4 family of cyclin-dependent kinase inhibitors directly enhance DNA damage repair independent of their role in cell cycle regulation (Cánepa et al., 2007). *CDKN2A* (p16^INK4a^), for example, protects cells from UV-induced apoptosis by downregulating the pro-apoptotic BAX protein (Al-Mohanna et al., 2004). After DNA damage, *CDKN1A* (p21^Cip1^) or *CDKN2A* overexpression induces cell cycle arrest and inhibits DNA damage induced apoptotic events such as cytochrome c release, mitochondrial membrane depolarization, and activation of the caspase cascade (Le et al., 2005). *CDKN2D* (p19^INK4d^) also plays a crucial role in regulating genomic stability and cell viability in response to genotoxic stress (Ceruti et al., 2005; Scassa et al., 2007; Tavera-Mendoza et al., 2006; Tavera-Mendoza et al., 2006). *CDKN2D* overexpression, for example, enhanced DNA damage repair whereas *CDKN2D*-deficient cells had diminished ability to repair DNA damage; Enhancement of DNA repair inversely correlated with apoptosis and was independent of its role in cell cycle regulation (Cánepa et al., 2007; Ceruti et al., 2005; Scassa et al., 2007).

Our finding that *CDKN2CRTG* overexpression protected Chinese hamster ovary cells from doxorubicin induced apoptosis likely extends the observation that the INK4 family of cyclin-dependent kinase inhibitors can enhance DNA damage repair to *CDKN2C*. While we have not explored the mechanisms by which *CDKN2CRTG* overexpression protects from apoptosis induced cell death, it is may be by slowing the progression of the cell cycle at the G_1_ and G_1_/S checkpoints and preventing rapid progression into G_2_, allowing cells more time to repair DNA damage or mechanisms independent of its role in cell cycle regulation like *CDKN2D*. *CDKN2CRTG* overexpression may also slow cell cycle progression into G_2_ when doxorubicin induces most DNA damage (Potter et al., 2002), thereby limiting the level of DNA damage and/or allowing cells to repair damage before DNA damage can initiate apoptosis. Thus, a longer cell cycle might allow Bowhead whale cells more time to correct DNA damage and other genomic instabilities such as aneuploidy but additional studies are needed to determine if the mechanism is direct, i.e., by allowing cells more time to repair DNA damage, indirect, i.e., by limiting the accumulation of DNA damage, or some combination of direct and indirect effects. Both indirect and direct effects models are consistent with observations that whale cells generally have more efficient DNA damage repair mechanisms than cells from some other mammals (Browning et al., 2017; Chen et al., 2009; Meaza et al., 2020; Wise et al., 2010, 2008).

### What might be the consequences of *CDKN2C* duplication? Reduced cancer risk

*CDKN2C* is a tumor suppressor for a variety of cancer types (Franklin et al., 2000), which may be related to both its role in cell cycle regulation and a direct role DNA damage repair, and this function is dosage sensitive. *CDKN2C* has a dosage dependent inhibitory effect on teratoma formation in NOD/SCID mice, such that tumor volume from *CDKN2C* overexpressing teratomas was significantly reduced compared to control and *CDKN2C*^−/−^ teratomas. Similarly, *CDKN2C* overexpression in the nasopharyngeal carcinoma epithelial tumor cell line HONE-1 prevented tumor growth when implanted to a nude mouse xenograft model (Li 2021,Lin 2018). *CDKN2C* haploinsufficiency also sensitizes mice to carcinogen-induced tumorigenesis (Bai et al., 2003). Remarkably, direct administration of a cell-permeable CDKN2C protein can inhibit tumor growth by 86-98% in a HCT116 tumor xenograft model (Lim et al., 2012) and *CDKN2C* copy number loss was associated with enhanced metastasis, metastasis to more distant sites, and decreased overall survival in patients with medullary thyroid carcinoma (Grubbs et al., 2016). These data indicate that *CDKN2C* dosage has functional effects on cancer initiation and progression and suggests that an additional copy of *CDKN2C* in Bowhead whales might be protective against the kinds of unregulated cell proliferation and genotoxic stress associated with cancer, particularly metastasis and survival once cancer develops.

### What might be the consequences of *CDKN2C* duplication? An aging, cancer resistance, and reproduction life history trade-off

While *CDKN2C* duplication may have played a role in the evolution of a remarkably long lifespan in Bowhead whales, our survey of mammalian *CDKN2C* copy number variation suggest that *CDKN2C* duplication is likely to be uncommon and thus may be associated with antagonistic pleiotropic effects that impose a constraint (cost) on increased dosage. Life history trade-offs between body size, lifespan, cancer resistance, and other organismal traits may be common (Garland, 2014; Stearns, 1989), particularly for reproductive (Boddy et al., 2015; Charnov and Ernest, 2006; Walker et al., 2008) and cellular phenotypes (Aktipis et al., 2013). For example, missense alleles in *ARHGAP27* are associated with increased number of children born but reduce reproductive lifespan (Mathieson et al., 2020), while missense alleles in *BRCA1* and *BRCA2* associated with increased lifetime risk of developing breast or ovarian cancer are also associated with an increased number of children born (Smith et al., 2012). Similarly, genetic variants that are associated with *HLA-F* and *TAP2* expression in humans increase fecundability but also predispose autoimmune disorders (Burrows et al., 2016; Mika et al., 2018; Mika and Lynch, 2016). Thus while additional copies of *CDKN2C* may provide cells with added robustness against the kinds of genotoxic stress associated with cancer progression, it may also be associated with antagonistic pleiotropic effects that constrain *CDKN2C* copy number changes.

*CDKN2C*, for example, is expressed in adipose tissue where it regulates adipocyte differentiation and lipid storage and *CDKN2C* expression levels have been associated with type-2 diabetes and central obesity (Pereira et al., 2022). Genome-wide association studies in the diversity outbred mouse population have also implicated *CDKN2C* in testis weight variability (Yuan et al., 2018). Consistent with this genetic association, *CDKN2C* is expressed in the seminiferous tubules of postnatal mice including post-mitotic spermatocytes undergoing meiosis (Yuan et al., 2018; Zindy et al., 2001) and *CDKN2C* knockout mice have generalized organomegaly including testicular enlargement and hyperplasia of interstitial testicular Leydig cells, which produce reduced levels of testosterone (Franklin et al., 2000, 1998; Zindy et al., 2001). While loss of *CDKN2C* (or *CDKN2D*) alone had no observable effect on male fertility, spermatogonia do not differentiate properly in *CDKN2C* and *CDKN2D* double-knockout mice and they produce few mature sperm; The residual spermatozoa have reduced motility and decreased viability (Yuan et al., 2018; Zindy et al., 2001). Thus, *CDKN2C* (*CDKN2D*) dosage is essential for testis and sperm development and male fertility.

In contrast to the testis phenotypes in *CDKN2C* in knockout mice, Bowhead whales have relatively small testes that are much smaller than than Right whales (~200kg vs >1,000kg), and “cryptic” Leydig (interstitial) cells that are difficult to identify by histological examination (O’Hara et al., 2002). Remarkably, several male pseudohermaphrodite Bowheads have also been described (O’Hara et al., 2002; Tarpley et al., 1995). While the external phenotype male pseudohermaphrodite Bowheads was female, the gonads were underdeveloped testes (O’Hara et al., 2002; Tarpley et al., 1995). In the most thoroughly studied individual, all derivatives of the mesonephric (Wolffian) and paramesonephric (Müllerian) ducts, the embryonic structures that form the internal male and female reproductive tract respectively, were absent (Tarpley et al., 1995). Male Bowhead whales also may have the oldest age at sexual maturation of any Cetacean (George et al., 1999). These phenotypes are consistent with Bowhead whales having an additional copy of *CDKN2C* and suggest tradeoff between aging and cancer resistance and male reproduction.

### Ideas and speculations: On the rapid evolution of cancer resistance

The increased longevity of Bowhead evolved very rapidly, within a few million years after their divergence with Right whales, which likely limited the kinds of genetic variation that could contribute to enhanced lifespan. For example, the average mammalian gene duplication rate, both segmental and retroduplication, is estimated to be 0.5–20×10^−3^ per gene per million years (Lynch and Conery, 2000; Pan and Zhang, 2007), suggesting that there should be about one gene duplication unique to Bowhead whales given Bowhead and Right whales diverged only ~4-5 million years ago (Árnason et al., 2018; Dornburg et al., 2012; McGowen et al., 2009). These data indicate that gene duplication has likely played a minor, even if important, role in the evolution of long lifespans and reduced cancer risk in Bowhead whales unlike the pervasive role gene duplication appears to have played in the evolution of reduced cancer risk in elephants, Xenarthrans, and Galapagos tortoises (Glaberman et al., 2021; Vazquez et al., 2022; Vazquez and Lynch, 2021). Thus, other mechanisms must have played a more numerous role than copy number gains and losses, such as gene regulatory evolution.

The evolutionary possibilities for solutions to Peto’s Paradox to arise is likely also constrained not just by the short time during which Bowheads evolved long lifesans, but also by the fundamental nature of developmental processes such as embryogenesis and growth rates that require cellular proliferation. We have shown that by insertion into a LINE1 element with a strong promoter, *CDKN2CRTG* achieves an extraordinarily high level of expression. This simple event likely led to an immediate phenotypic consequence in individuals with the *CDKN2CRTG* duplication, and possibly immediate reduction in cancer risk, that may have facilitated any ongoing increases in body size and lifespan in the Bowhead Lineage. Of course, to prove such a hypothesis requires the discovery of additional Mystecini fossils to reconstruct the history of body size expansions within this clade, which could then be correlated with the gene duplications described herein with functional validation; A developmental and evolutionary proof of the importance of the *CDKN2CRTG* duplication for body size and lifespan evolution in Bowheads would also require the kinds forward and reverse genetic experiments that are possible in model organisms but impossible in whales.

### Caveats and limitations

Our gain of function experiments indicate that *CDKN2CRTG* expression may affect the cell cycle length and the response of cells to DNA damage. A limitation of these experiments, however, is that we have not directly quantified DNA damage and repair, therefore it is unclear if the protective effects of *CDKN2CRTG* overexpression we observed is because of direct effects on DNA damage and repair rates or indirect effects based on cell cycle slowing. To disentangle these possibilities, we must quantify DNA damage repair rates between cells in different cycle phases and with *CDKN2CRTG* overexpression. We must also perform similar gain and loss of function experiments in Bowhead whale cells to establish a causal connection between *CDKN2CRTG* and cellular phenotypes related to healthy aging and lifetime cancer resistance.

Finally, we are unable to determine causality, i.e. that a specific genetic change, such as the retroduplication of *CDKN2C*, is necessary and sufficient for the realization of phenotypes that contribute to the evolution of long lifespans in Bowhead whales using traditional forward and reverse genetic techniques. While this constrains our ability to demonstrate proximate causation within an organismal context, we can construct strong circumstantial arguments in favor of causation using methods like those used to associate genetic variation with phenotypes in genome-wide association studies.

## Conclusions

Bowhead whales are an excellent model system in which to explore the evolution of long lifespans because they are closely related to species with much shorter lifespans and evolved that long lifespan since their divergence with Right whales ~4-5 million years ago. In this study we used comparative genomics to identify gene duplications that are phylogenetically associated with the evolution of long lifespans in Bowhead whales. We identified a single gene duplication, a retroduplication of the cyclin dependent kinase inhibitor *CDKN2C* (*CDKN2CRTG)*, unique to Bowhead whales. Remarkably, the putative phenotypic consequences of a duplicate *CDKN2C* gene are relatively straightforward given its role in cell cycle regulation and DNA repair. Our experimental data indicate that *CDKN2CRTG* may regulate the cell cycle and contribute to enhanced DNA damage repair. These data suggest that Bowhead whales evolved their extremely long lifespans at least in part through duplication of the *CDKN2C* gene, which may reduce their lifetime risk of developing cancer.

## Materials and methods

### Identification of gene duplications in the Bowhead, Right, and Sperm whale genomes

#### Approach

We identified putative gene duplicates in the genome of a Bowhead whale (PRJNA194091), North Atlantic (https://www.dnazoo.org/assemblies/Eubalaena_glacialis) and South Atlantic (https://www.dnazoo.org/assemblies/Eubalaena_australis) Right whales, and Sperm whale (assembly ASM283717v2) using a combination of Reciprocal Best Hit BLAT (RBHB) and Effective Copy Number By Read Coverage (ECNC), which we have previously described (Vazquez and Lynch, 2021). Note that the draft Right whale assemblies was generated by the DNA Zoo team and Mark Yandell’s lab at the University of Utah with contributions from Michael S. Campbell, Brian Dalley, Edgar J. Hernandez, Barry Moore, Andrea Chirife, Matias Di Martino, Mariano Sironi, Luciano O. Valenzuela, Marcela Uhart, Victoria J. Rowntree, Guy D. Eroh, Sancy A. Leachman and Jon Seger. Construction of the draft was funded and supported by the Illumina Corporation, by the H.A. and Edna Benning Fund, and by the US National Science Foundation.

#### Reciprocal best hit BLAT

We previously developed a reciprocal best hit BLAT pipeline to identify putative homologs and estimate gene copy number across species (Vazquez and Lynch, 2021). The Reciprocal Best Hit (RBH) search strategy is conceptually straightforward: 1) Given a gene of interest *G_A_* in a query genome *AA*, one searches a target genome *B* for all possible matches to *G_A_ G*_A_; 2) For each of these hits, one then performs the reciprocal search in the original query genome to identify the highest-scoring hit; 3) A hit in genome *B* is defined as a homolog of gene *G_A_* if and only if the original gene *G_A_ G*_A_ is the top reciprocal search hit in genome *AA*. We selected BLAT as our algorithm of choice, as this algorithm is sensitive to highly similar (>90% identity) sequences, thus identifying the highest-confidence homologs while minimizing many-to-one mapping problems when searching for multiple genes. RBH performs similar to other more complex methods of orthology prediction and is particularly good at identifying incomplete genes that may be fragmented in low quality/poorly assembled regions of the genome.

#### Estimated copy number by coverage

In low-quality genomes, many genes are fragmented across multiple scaffolds, which results in BLA(S)T-like methods calling multiple hits when in reality there is only one gene. To compensate for this, we developed a novel statistic, Estimated Copy Number by Coverage (ECNC), which averages the number of times we hit each nucleotide of a query sequence in a target genome over the total number of nucleotides of the query sequence found overall in each target genome (Vazquez and Lynch, 2021). This allows us to correct for genes that have been fragmented across incomplete genomes, while accounting for missing sequences from the human query in the target genome. Details of the ECNC method can be found in (Vazquez and Lynch, 2021).

#### RecSearch pipeline

We created a custom Python pipeline for automating RBHB searches between a single reference genome and multiple target genomes using a list of query sequences from the reference genome. For the query sequences in our search, we used the hg38 UniProt proteome, which is a comprehensive set of protein sequences curated from a combination of predicted and validated protein sequences generated by the UniProt Consortium. Next, we excluded genes from downstream analyses from large multigene families for which assignment of homology using the RBBH approach was uncertain, including uncharacterized ORFs, LOCs, HLA genes, replication dependent histones, odorant receptors, ribosomal proteins, zinc finger transcription factors, Coiled Coil Domain Containing proteins, Family With Sequence Similarity genes, Mitochondrial Ribosomal Proteins, NADH:Ubiquinone Oxidoreductase, Solute Carrier Family genes, TMEM Transmembrane Proteins, DnaJ Heat Shock Protein Family genes, Dual Specificity Phosphatases, Eukaryotic Translation Initiation Factors, F-Box Proteins, Polypeptide N-acetylgalactosaminyltransferases, Ring Finger Proteins, Sterile Alpha Motif Domain Containing proteins, Serpins, SEC14 And Spectrin Domain Containing proteins, Small Nuclear Ribonucleoproteins, Serine/Threonine Kinases, TBC1 Domain Family Members, Tubulins, Ubiquitin Conjugating Enzymes, WD domain containing proteins, viral and repetitive-element-associated proteins, “Uncharacterized,” “Putative,” and “Fragment” proteins.

#### Duplicate gene inclusion criteria

In order to condense transcript-level hits into single gene loci, and to resolve many-to-one genome mappings, we removed exons where transcripts from different genes overlapped, and merged overlapping transcripts of the same gene into a single gene locus call. The resulting gene-level copy number table was then combined with the maximum ECNC values observed for each gene in order to call gene duplications. We called a gene duplicated if its copy number was two or more, and if the maximum ECNC value of all the gene transcripts searched was 1.5 or greater; We have shown that incomplete duplications can encode functional genes (Sulak et al., 2016b; Vazquez et al., 2018), therefore partial gene duplications were included provided they passed additional inclusion criteria (see below). The ECNC cut-off of 1.5 was selected empirically, as this value minimized the number of false positives seen in a test set of genes and genomes. We thus identified 1710 duplicate genes in the Bowhead whale genome. We note that: 1) The RBBH/ECNC approach is not exhaustive and other gene duplications that are 100% identical between query and target sequences within the Bowhead whale genome may be collapsed into a single hit, undercounting gene duplicates; 2) These duplicate genes may not be specific to Bowhead whales and represent a mixture of both ancient and recent gene duplications; and 3) We have not filtered these duplicates to exclude those with signatures of pseudogenes such as inframe stop codons, retrogenes, or non-expressed genes. Thus this duplicate gene set will include a mixture of functional and non-functional genes, but is a useful starting point to more detailed studies such as manual verification of duplicate genes.

### Identification of the *CDKN2C* gene in additional Sarcopterygian and Cetacean genomes

We also used BLAT to verify the copy number of CDKN2C in 61 Sarcopterygian genomes (**Figure 1 – source data 2**). After identifying the canonical gene from each species, we used the corresponding nucleotide sequences as the query sequence for additional BLAT searches within that species’ genome. To further confirm the orthology of each gene we used a reciprocal best BLAT approach, sequentially using the putative CDS of each gene as a query against the human genome.

### RNA-Seq transcript abundance estimation

We used previously published RNA-Seq data from seven Bowhead whale tissues (Keane et al., 2015); five Minke whale tissues (Yim et al., 2014); one Humpback whale tissue; one Sperm whale tissue; and data from 454 GS FLX pyrosequencing of five dolphin tissues (Westbury et al., 2019) to estimate transcript abundances (**Figure 3 – source data 1**). Individual datasets were aligned to the Bowhead whale, Minke whale, Humpback whale, Sperm whale, or dolphin reference genome on Galaxy with HISAT2 (Galaxy Version 2.2.1+galaxy0) using default settings and reporting alignments tailored for transcript assemblers including StringTie, and transcripts assembled with StringTie (Galaxy Version 2.1.7+galaxy1) without a reference to guide assembly. Assembled transcripts from each dataset were merged into a single gtf file with StringTie merge transcripts (Galaxy Version 2.1.7+galaxy1) using default settings, except that minimum input transcript abundances (FPKM and TPM) to include in the merge was set to 0. Sequences for each transcript were extracted from the merged gtf file with gffread (Galaxy Version 2.2.1.3+galaxy0) and used as the reference transcriptome to estimate transcript abundances with Kallisto and bias correction (Bray et al., 2016).

### Fucci2a construct synthesis and stable cell line generation

We generated a Fucci2a reporter vector by synthesizing the mCherry-hCdt1(30/120) (GenBank accession: AB512478), *Thosea asigna* virus 2A (T2A) self-cleaving peptide (GSGEGRGSLLTCGDVEENPGP), and mVenus-hGeminin(1/110) (GenBank accession: AB512479) reporters as a single gene with mouse codon usage (GeneScript);. The mCherry-hCdt1(30/120)-T2A-mVenus-hGeminin(1/110) fusion gene was then cloned into the pcDNA5/FRT vector (GeneScript). We generated a stable Fucci-2a reporter Chinese Hamster Ovary (FuCHO) cell line using the Flp-In System (Thermo Fisher K601001) and the Flp-In-CHO cell line (invitrogen R758-07)). Flp-In-CHO Chinese Hamster ovary cells were maintained in T75 culture flasks at 37°C, 5% CO2, and 90-90% humidity in a culture medium consisting of Ham’s F12, 10% FBS, 2mM L-glutamine, 1% Pen-Strep, and 100ug/mL of Zeocin (to maintain stable integration of the FRT landing site). To integrate the Fucci-2a reporter into the Flp-In-CHO cell line, 1×10^6^ cells were transiently with the Fucci-2a pcDNA5/FRT vector using Lipofectamine LTX (Invitrogen), the CHO-K1 transfection protocol, and the recommended ratio of insert plasmid and flippase plasmid. 24 hours after transfection the cells were selected using 800ug/mL hygromycin (Gibco) for 5 days. Cells were propagated in media containing 200ug/mL Hygromycin. Stability of the knock-in was confirmed by estimating the abundance of fluorescent cells in culture over several passages, both of the original lines as well as from cryopreserved aliquots.

### Luciferase reporter vector construction and Dual-Luciferase Reporter assay

To test if the region 5’ to the *CDKN2CRTG* gene regulatory abilities, we divided it into ten 301bp fragments including 51bp 5’- and 3’-overlaps that tile the region from upstream of the L1-2_Ttr element to downstream of the *CDKN2CRTG* initiation codon using OVERFRAG (http://143.107.58.249/cgi-bin/overfrag.cgi) (Ferreira-Junior and Digiampietri, 2019). Each fragment was synthesized and cloned (GeneScript) into the pGL4.14[*luc2*/Hygro] and pGL4.26[*luc2*/minP/Hygro] firefly luciferase reporter vectors (Promega); the pGL4.14[*luc2*/Hygro] vector lacks an enhancer and a promoter whereas the pGL4.26[*luc2*/minP/Hygro] vector contains a minimal promoter. Luciferase assays were performed in FuCHO cells, which were maintained in T75 culture flasks at 37°C, 5% CO2, and 90-98% humidity in a culture medium consisting of Ham’s F12, 10% FBS, 2mM L-glutamine, 200ug/mL Hygromycin, and 1% Pen-Strep. 10^4^ cells were harvested and seeded into 96-well white culture plates. 24 hours later, cells were transfected using Lipofectamine LTX (Invitrogen) and either 100ug of the pGL4.14[*luc2*/Hygro] or pGL4.26[*luc2*/minP/Hygro] vectors and 1ng of the pGL4.74[hRluc/TK] *Renilla* control reporter vector according the standard protocol, with 0.5 mL/well of Lipofectamine LTX reagent and 0.1 mL/well of PLUS reagent. Luminescence was quantified 48 hours after transfection using a Dual-Luciferase Reporter Assay System (Promega) in a GloMax-Multi+ Reader (Promega). For all experiments, firefly luminescence was standardized to *Renilla* luminescence to control for differences in transfection efficiency across samples, and firefly/*Renilla* luminescence values of firefly luciferase vectors with tiling inserts were standardized to firefly/*Renilla* luminescence values of firefly luciferase vectors without tiling inserts.

### CDKN2CRTG-eGFP construct synthesis and flow cytometry methods

We synthesized the Bowhead whale *CDKN2CRTG* protein with mouse codon usage (GeneScript) and directly cloned the synthesized gene into the pcDNA3.1-C-eGFP vector (Life Technologies), which added a C-terminal eGFP tag. Cell cycle analyses were performed in FuCHO cells, which were maintained in T75 culture flasks as described. FUCHO cells were transfected with either the CDKN2CRTG-eGFP transgene plasmid, a GFP-control plasmid, or a DNA-free mock transfection. Cells were grown for 48 hours as described in the prior section. Furthermore, FLP-IN CHO cells were transfected with either a mock solution or the eGFP plasmid as a no-color and GFP controls. Cells were harvested using 0.05% Trypsin-EDTA and washed three times with DPBS. Cells were resuspended at a concentration of 1 million cells/mL in DPBS, and then fixed in 1% paraformadehyde for 15 minutes at room temperature. Cells were then washed twice with DPBS and resuspended at a concentration of 1-2 million cells/mL in 10% BSA in DPBS for flow cytometry. Cells were analyzed using a LSR Fortessa (BD Biosciences) configured with 488nm, 640nm, 561nm and 405nm lasers using FCS Express. Due to the high transfection efficiency, the eGFP sample was spiked with 10% mock transfected FLP-IN CHO control as an internal negative control. Voltages, gates, and compensation for eGFP (520/50), mCherry (610/20), and mVenus (520/50) were calibrated using the eGFP-only and mock transfected FuCHO cells. 10,000 events were captured for each sample after calibration. All analyses were done using the FACS Diva software.

### ApoTox-Glo Triplex Assay

ApoTox-Glo Triplex Assays (Promega) were performed in FuCHO cells, which were maintained in T75 culture flasks at 37°C, 5% CO2, and 90-90% humidity in a culture medium consisting of Ham’s F12, 10% FBS, 2mM L-glutamine, and 1% Pen-Strep. 5×10^4^ cells were harvested and seeded into 96-well white culture plates and reverse-transfected with 100ng of the CDKN2CRTG-eGFP reporter vector using Lipofectamine LTX (Invitrogen) according the standard protocol, with 0.5 mL/well of Lipofectamine LTX reagent and 0.1 mL/well of PLUS reagent. 24 hours after transfection cells were treated with 0, 0.5, 1, 5, 10, or 50uM doxorubicin for 6 or 18 hours. The ApoTox-Glo Triplex Assay (Promega) was performed using the default ApoTox-Glo Triplex Assay in a GloMax-Multi+ Reader (Promega).

### Mammalian population doubling time estimation

We gathered population doubling times from previously published studies (Burkard et al., 2019, 2015; Carvan et al., 1994; Chen et al., 2009; Gomes et al., 2011; Mancia et al., 2012; Rajput et al., 2018; Seluanov et al., 2008; Wang et al., 2011, 2021; Wise et al., 2011, 2008; Yajing et al., 2018). Some studies did not report doubling times, but showed graphs of doubling times. For these datasets, doubling times were extracted from figures using WebPlotDigitizer Version 4.5 (Rohatgi, 2021).

### Bowhead whale cell culture and doubling time estimation

Bowhead whale primary kidney fibroblast cells were maintained in T75 culture flasks at 37°C/5% CO2 in a culture medium consisting of fibroblast growth medium (FGM)/Eagle’s minimum essential medium (EMEM) (1:1) supplemented with insulin, fibroblast growth factor (FGF), 6% FBS, and gentamicin/amphotericin B (FGM-2; singlequots; Clonetics/Lonza). At 80% confluency, cells were harvested and seeded into six-well culture plates at 10,000 cells/well. Once cells recovered to 80% confluency, they were harvested by trypsinization. Cells were manually counted using trypan blue staining and a hemocytometer.

### Cancer data collection from previous studies

Published cancer prevalence data for *Delphinapterus leucas (Martineau et al., 2002, 1988)*, *Globicephala melaena (Albuquerque et al., 2018)*, *Tursiops truncatus (Albuquerque et al., 2018)*, *Phocoena phocoena (Albuquerque et al., 2018)*, *Balaena mysticetus (R. et al., 2020; Stimmelmayr et al., 2017)*, *Eubalaena glacialis (Sharp et al., 2019)*, *Ziphius cavirostris* (Rogan et al., 1997) was collected from published surveys of morbidity and mortality in these species. In addition, we also collected data from several studies reporting cancer prevalence aggregated across multiple species (Díaz-Delgado et al., 2018; Geraci et al., 1987; Uys and Best, 1966). Neoplasia prevalence in these Cetaceans were compared to to other Therian mammals using data from two published studies that included pathology reports from 37 (Boddy et al., 2020) and 191 (Vincze et al., 2022) species of Therian mammals, as well as several species of Xenarthrans we also collected from the literature, including giant anteater (*Myrmecophaga tridactyla*), southern tamandua (*Tamandua tetradactyla*), silky anteater (*Cyclopes* sp.), maned three-toed sloths (*Bradypus torquatus*), brown-throated sloths (*Bradypus variegatus*), pale-throated sloth (*Bradypus tridactylus*), Linné’s two-toed sloth (*Choloepus didactylus*), three-toed sloths (*Bradypus sp*.), and two-toed slots (*Choloepus sp*.) (Arenales et al., 2020a, 2020b; Diniz and Oliveira, 1999; Diniz et al., 1995); The total dataset includes cancer prevalence data from 221 species. Neoplasia reports from *Dasypus novemcinctus* from Boddy et al. were combined with cancer prevalence data from the NHDP colony (Vazquez et al., 2022) and *Choloepus didactylus* data from Vincze et al were combined with *Choloepus didactylus* data from (Arenales et al., 2020b). Confidence intervals (95%) on lifetime neoplasia prevalence were estimated in PropCI package (https://github.com/shearer/PropCIs) in R using the Clopper-Pearson exact CI function: exactci(x, n, conf.level=0.95), where x is the number of successes (necropsies with neoplasia), n is the total sample size (number of necropsies), and conf.level=0.95 is the 95% lower and upper confidence intervals.

### Data analysis

All data analysis was performed using Python version 3.8 and R version 4.0.2 (2020-06-22). Scripts for performing the reciprocal best hit BLAT and ECNC correction were described in Vazquez and Lynch (2021, https://github.com/docmanny/atlantogenataGeneDuplication) copy archived at swh:1:rev:6bc68ac31ef148131480710e50b0b75d06077db2. All files necessary to reproduce the data in this manuscript are provided in Source data 1.

### Cell lines

Flp-In-CHO Chinese Hamster ovary cells, which are derived from the CHO-K1 cell line (ATCC CCL-61), were purchased from ThermoFisher (invitrogen R758-07); their identity has been authenticated by Invitrogen. Bowhead whale primary kidney fibroblast cells were provided by the T.C. Hsu Cryo-Zoo at the University of Texas MD Anderson Cancer Center. Both cell lines were determined to be mycoplasma free.

## Data availability

All genome data analyzed during this study are publicly available (**Figure 1 - source data 1**) all other data generated or analyzed in this study are included in the manuscript and supporting files. A supplemental R markdown file is provided that includes raw data and scripts to reproduce the luciferase assay data shown in Figure 3, Figure 3 – figure supplement 1, and Figure 3 – figure supplement 2, the flow data shown in Figure 4, and the ApoTox-Glo data shown Figure 5.

## Acknowledgements

We thank R. Behringer and the T.C. Hsu Cryo-Zoo (University of Texas MD Anderson Cancer Center) for providing the Bowhead whale primary kidney fibroblast cell line.

---

**Figure 4 – figure supplement 1.**
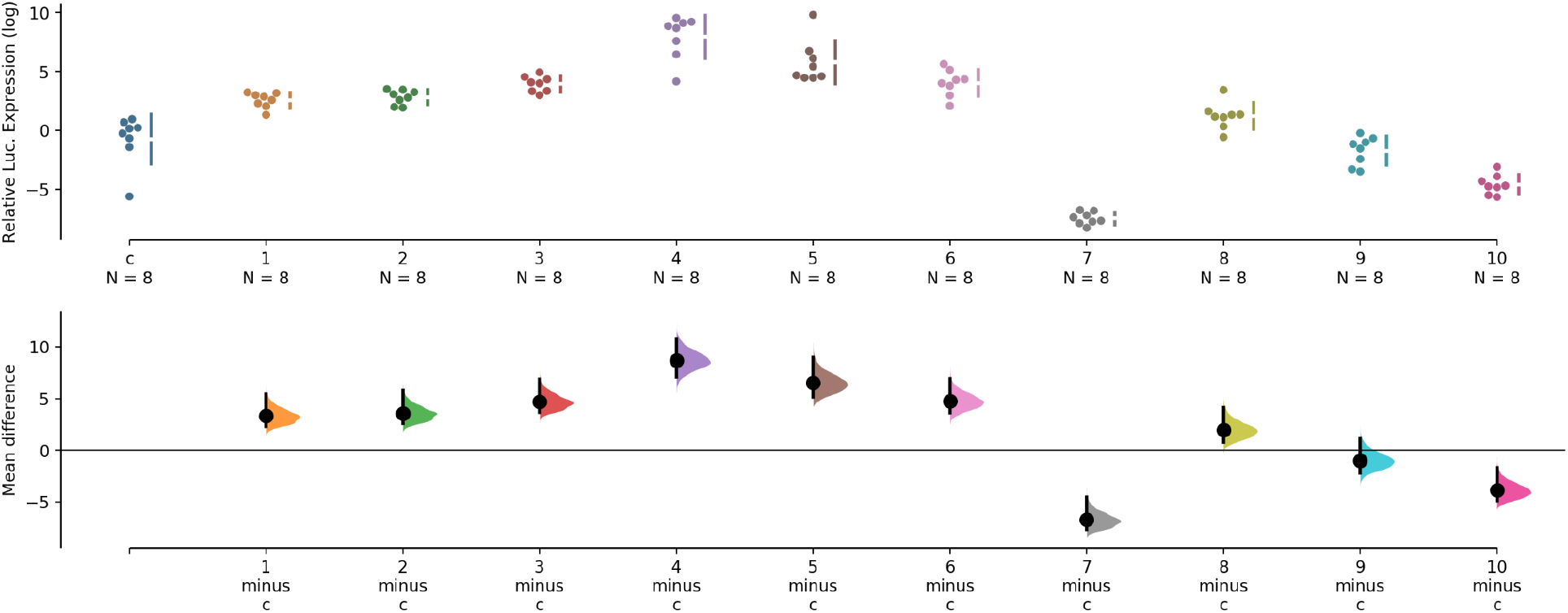
pGL4.14[*luc2*/Hygro] dual luciferase reporter assay results. Ten 301bp overlapping fragments that tile the region from upstream of the L1-2_Ttr element to downstream of the *CDKN2CRTG* initiation codon were cloned into luciferase reporter vector pGL4.14[*luc2*/Hygro], which lacks both an enhancer and a promoter, and in transiently transfected mouse 3T3 cells. Relative luciferase expression (standardized to *Renilla* and empty vector control), n=8. The mean difference for 10 comparisons against the shared empty vector control (c) are shown as a Cumming estimation plot. The raw data is plotted on the upper axes. On the lower axes, mean differences are plotted as bootstrap sampling distributions. Each mean difference is depicted as a dot. Each 95% confidence interval is indicated by the ends of the vertical error bars. The effect sizes and CIs are reported above as: effect size [CI width lower bound; upper bound]. 5000 bootstrap samples were taken; the confidence interval is bias-corrected and accelerated. The P value(s) reported are the likelihood(s) of observing the effect size(s), if the null hypothesis of zero difference is true. Results: The unpaired mean difference between c and fragment 1 is 3.31 [95.0%CI 2.31, 5.43]. The P value of the two-sided permutation t-test is 0.0. The unpaired mean difference between c and fragment 2 is 3.57 [95.0%CI 2.57, 5.77]. The P value of the two-sided permutation t-test is 0.0. The unpaired mean difference between c and fragment 3 is 4.69 [95.0%CI 3.69, 6.89]. The P value of the two-sided permutation t-test is 0.0. The unpaired mean difference between c and fragment 4 is 8.7 [95.0%CI 7.07, 10.7]. The P value of the two-sided permutation t-test is 0.0. The unpaired mean difference between c and fragment 5 is 6.53 [95.0%CI 5.16, 9.0]. The P value of the two-sided permutation t-test is 0.0002. The unpaired mean difference between c and fragment 6 is 4.77 [95.0%CI 3.58, 6.89]. The P value of the two-sided permutation t-test is 0.0. The unpaired mean difference between c and fragment 7 is −6.73 [95.0%CI −7.7, −4.55]. The P value of the two-sided permutation t-test is 0.0. The unpaired mean difference between c and fragment 8 is 1.96 [95.0%CI 0.779, 4.15]. The P value of the two-sided permutation t-test is 0.0116. The unpaired mean difference between c and fragment 9 is −0.991 [95.0%CI −2.2, 1.09]. The P value of the two-sided permutation t-test is 0.257. The unpaired mean difference between c and fragment 10 is −3.86 [95.0%CI −4.91, −1.7]. The P value of the two-sided permutation t-test is 0.0004.

**Figure 4 – figure supplement 2.**
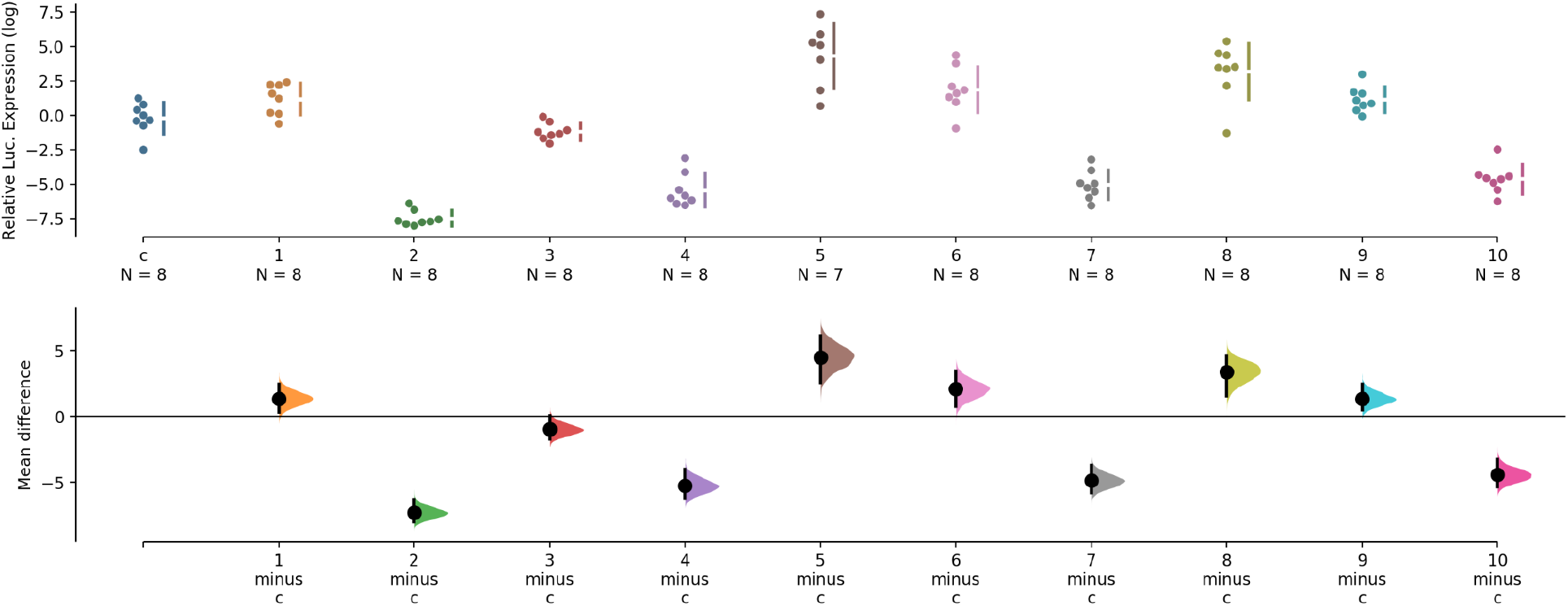
pGL4.26[*luc2*/minP/Hygro] dual luciferase reporter assay results. Ten 301bp overlapping fragments that tile the region from upstream of the L1-2_Ttr element to downstream of the *CDKN2CRTG* initiation codon were cloned into luciferase reporter vector pGL4.26[*luc2*/minP/Hygro], which lacks an enhancer but has a minimal promoter, and in transiently transfected mouse 3T3 cells. Relative luciferase expression (standardized to *Renilla* and empty vector control), n=8. The mean difference for 10 comparisons against the shared empty vector control (c) are shown as a Cumming estimation plot. The raw data is plotted on the upper axes. On the lower axes, mean differences are plotted as bootstrap sampling distributions. Each mean difference is depicted as a dot. Each 95% confidence interval is indicated by the ends of the vertical error bars. The effect sizes and CIs are reported above as: effect size [CI width lower bound; upper bound]. 5000 bootstrap samples were taken; the confidence interval is bias-corrected and accelerated. The P value(s) reported are the likelihood(s) of observing the effect size(s), if the null hypothesis of zero difference is true. Results: The unpaired mean difference between c and 1 is 1.36 [95.0%CI 0.326, 2.45]. The P value of the two-sided permutation t-test is 0.0292. The unpaired mean difference between c and 2 is −7.29 [95.0%CI −8.0, −6.32]. The P value of the two-sided permutation t-test is 0.0. The unpaired mean difference between c and 3 is −0.972 [95.0%CI −1.7, 0.042]. The P value of the two-sided permutation t-test is 0.0462. The unpaired mean difference between c and 4 is −5.26 [95.0%CI −6.19, −4.01]. The P value of the two-sided permutation t-test is 0.0. The unpaired mean difference between c and 5 is 4.5 [95.0%CI 2.6, 6.12]. The P value of the two-sided permutation t-test is 0.0002. The unpaired mean difference between c and 6 is 2.08 [95.0%CI 0.749, 3.44]. The P value of the two-sided permutation t-test is 0.013. The unpaired mean difference between c and 7 is −4.86 [95.0%CI −5.78, −3.69]. The P value of the two-sided permutation t-test is 0.0. The unpaired mean difference between c and 8 is 3.38 [95.0%CI 1.55, 4.59]. The P value of the two-sided permutation t-test is The unpaired mean difference between c and 9 is 1.35 [95.0%CI 0.504, 2.44]. The P value of the two-sided permutation t-test is 0.0168. The unpaired mean difference between c and 10 is −4.43 [95.0%CI −5.32, −3.25]. The P value of the two-sided permutation t-test is 0.0002.

**Figure 5 – figure supplement 1.**
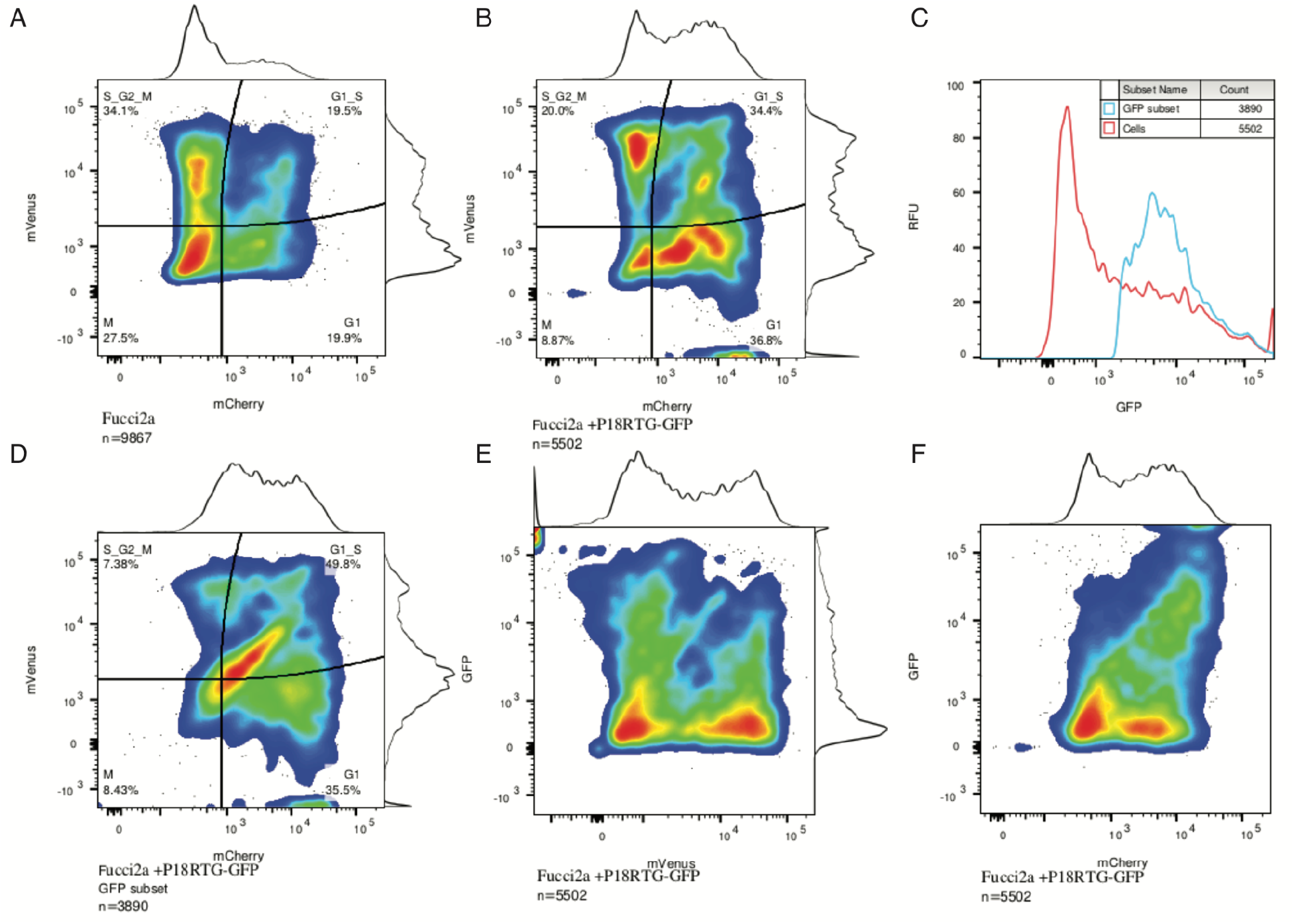
**(A)** The distribution of cell cycle states in mock transfected Fucci2a CHO cells, 19.5% of cells are in G1/S. **(B)** The distribution of cell cycle states in *CDKN2CRTG* transfected Fucci2a CHO cells. Note the shift in cells towards G1 and G1/S phase from 19.5% to 34.4%. **(C)** Gated subset of GFP-cells shown in F. **(D)** The distribution of cell cycle states GFP gated *CDKN2CRTG* transfected Fucci2a CHO cells. Note the shift in cells towards G1 and G1/S phase from 7.38% to 49.8%. The shift in the population to G1 and especially G1/S is clear. **(E)** GFP levels as a function of mVenus in the ungated populations. **(F)** GFP levels as a function of mCherry in the ungated populations.

